# A Wireless Optoelectronic Probe Monitors Tissue Oxygenation in the Deep Brain

**DOI:** 10.1101/2023.05.21.541646

**Authors:** Xue Cai, Haijian Zhang, Penghu Wei, Quanlei Liu, Dawid Sheng, Zhen Li, Wenxin Zhao, Zhongyin Ye, Zhao Xue, Yang Xie, Yang Dai, Changming Wang, Yuqi Wang, Xin Fu, Bozhen Zhang, Lan Yin, Hongshang Peng, He Ding, Guoguang Zhao, Xing Sheng

## Abstract

Real-time detection of tissue oxygenation in the nervous system is crucial for neuroscience exploration and clinical diagnostics. Compared to blood oxygenation, the partial pressure of oxygen in brain tissue (PbtO_2_) possesses more direct relevance to local neural activities and metabolic conditions. In this paper, we present an implantable optoelectronic probe that wirelessly and continuously monitors PbtO_2_ signals in the deep brain of living animals. The thin-film, microscale implant integrates a light-emitting diode and a photodetector coated with oxygen sensitive dyes. Powered by a battery or an inductive coil, a miniaturized circuit is capable of recording and wirelessly transmitting PbtO_2_ signals, which allows for simultaneous monitoring of PbtO_2_ levels in multiple freely moving rodents. The wireless micro-probe captures cerebral hypoxia states of mice in various scenarios, including altered inspired oxygen concentration, acute ischemia. Particularly, in mouse models with seizures, the micro-probe associates temporal PbtO_2_ variations in multiple brain regions with electrical stimulations imposed in the hippocampus. These materials and device strategies overcome the limits of existing oxygen sensing approaches and provide important insights into neurometabolic coupling.

## INTRODUCTION

Oxygen supply is vital in human metabolism. Hypoxia, which reveals that tissue oxygen is below the normal level, can cause dysfunction, damage and even death of biological cells and organs^1–3^. As a leading consumer of oxygen, the human brain, which constitutes only ∼2% of the body weight, expends ∼20% of the body’s total oxygen supply to continuously maintain its normal function^4^. Caused by acute brain abnormalities such as poisoning, traumatic brain injury and subarachnoid hemorrhage^5–7^, or diseases including hydrocephalus and brain tumors, ^8, 9^ ^10^, cerebral hypoxia can lead to irreversible damage in brain functions and result in disability and death. Additionally, cerebral hypoxia is closely associated with neurological and cerebrovascular diseases like epilepsy and apoplexy, as well as various neurodegenerative disorders^11, 12, 13^. Besides cell survival, oxygen also modulates cell proliferation and differentiation and influences neurogenesis^14^. Therefore, real-time monitoring the cerebral oxygenation state is crucial not only in understanding the metabolic coupling between oxygen and brain activities, but also for improving clinical practices^15–17^.

Conventional biological techniques to detect oxygen (O_2_) in the brain like near infrared spectroscopy (NIRS) and functional magnetic resonance imaging (fMRI) are limited by data inaccuracies and low spatial-temporal resolutions^18–20^. Furthermore, these non-invasive techniques, along with other recently reported invasive O_2_ sensing probes^21, 22^, only assess the cerebral blood oxygen saturation by exploiting the difference of optical absorption (for NIRS) or magnetic (for fMRI) properties between oxygenated and deoxygenated hemoglobin. While tissue oxygen saturation (StO_2_) reflects the level of oxygen in the blood transported throughout the body, its value sometimes deviates from the partial pressure of oxygen in brain tissue (PbtO_2_), which directly represents the amount of oxygen delivered to and consumed by cells and tissues^23^. Non-invasive methods for the direct PbtO_2_ detection involve fluorine-based MRI^24^, electron paramagnetic resonance (EPR) ^25^ and positron emission tomography (PET) ^26^. With millimeter-scale resolution, these measurements rely on injected or inhaled oxygen tracers that delay the PbtO_2_ detection, and bulky, high-cost instruments that prohibit portable use.

To measure local PbtO_2_ signals in the deep brain, clinically available techniques utilize the insertion of electrochemical or optical probes into the targeted region. Electrochemical methods based on polarographic analyses employ Clark-type electrodes or other conductive wires to detect PbtO_2_ signals via oxidative reactions ^27–30^. There are also reports on wirelessly operated electrochemical sensors used in rodents^31^. These chemical electrodes typically consume oxygen during detection, take a certain time to stabilize, and are vulnerable to contamination from interfering substances in the absence of a carefully designed O_2_ selective membrane^31, 32^. Optical techniques leverage phosphorescent dyes whose luminescence efficiencies or lifetimes decay with the addition of oxygen^33^, so that the tissue oxygenation can be sensed by imaging or spectroscopic setups^34–36^. Compared with electrochemical probes, the luminescence quenching process exhibits superior oxygen selectivity and promising resistance to electromagnetic interferences^37, 38^. Traditional tools based on this mechanism to sense PbtO_2_ utilize tethered fiber-optic systems for light delivery and collection within the deep tissue^39^. Wirelessly operated optical implants, which directly monitor real-time PbtO_2_ signals in freely moving animals, are highly desirable. A recent study remarkably demonstrates optical-based oxygen sensing in the deep peripheral tissue in living sheep with ultrasound-powered, miniaturized optoelectronic devices^40^; however, its deployment in the deep brain has not yet been accomplished, probably constrained by the limited ultrasound penetration through the skull. Therefore, we anticipate that wireless optoelectronic implants with the capability for dynamic PbtO_2_ monitoring would provide unprecedented insights into brain function and metabolism, advance neuroscience research and improve healthcare.

In this paper, we present a microscale, optoelectronic probe that wirelessly monitors in vivo PbtO_2_ signals in real time. The thin-film micro-probe measures local PbtO_2_ levels based on luminescent quenching of phosphorescent dyes, whose optical signals are excited and collected by integrating a light-emitting diode (LED) and a photodetector. We implement miniaturized circuits, which can be powered by a battery or an inductive coil, to capture and wirelessly transmit the PbtO_2_ data. In vitro and in vivo studies reveal the system’s capability of dynamic PbtO_2_ recording. Implanted into the deep brain tissue, the micro-probe helps correlate cerebral hypoxia conditions with electrophysiological activities in a rodent seizure model. These device strategies promise previously inaccessible research in neuroscience studies and clinical applications.

## RESULTS

### The microscale optoelectronic probe for PbtO_2_ monitoring

We design and fabricate a microscale optoelectronic probe to monitor PbtO_2_ changes in the deep brain, schematically illustrated in Figure 1a. Figure 1b provides an exploded depiction of the micro-probe structure, and the detailed probe design and fabrication procedures are described in the Methods and Figure S1. From top to bottom, the micro probe consists of a polydimethylsiloxane (PDMS) film embedded with platinum(II) 5,10,15,20-tetrakis-(2,3,4,5,6-pentafluorphenyl)-porphyrin (PtTFPP) phosphors, a PDMS/parylene encapsulation layer, an indium gallium nitride (InGaN) violet LED, a dielectric filter, and an indium gallium phosphide (InGaP) photodetector on a polyimide (PI) substrate. The PbtO_2_ sensing mechanism operates on the principle of the photoluminescence (PL) quenching of PtTFPP, which exhibits enhanced PL when the surrounding O_2_ level drops^41^ (Figure 1c and 1d). PtTFPP with a concentration of 0.1 wt% is mixed into the PDMS coating layer with a thickness of 10 μm, to obtain optimal absorption and emission properties (Figure S2). The PtTFPP/PDMS film absorbs violet light (with the absorption peak at 389 nm) and shows red PL emission peaked at 646 nm (Figure 1e). With an electroluminescent (EL) peak wavelength of 395 nm, the InGaN violet LED serves as an excitation source for PtTFPP, whose PL emission is captured by the InGaP detector. In particular, the multilayered silicon oxide (SiO_2_)/titanium oxide (TiO_2_)-based longpass filter on the InGaP detector effectively optimizes its sensitivity in the red spectral range, resulting in a band-selective absorption between 570 nm and 655 nm. These thin-film, microscale devices, including the InGaN LED, the InGaP detector and the dielectric filter, are fabricated through epitaxial liftoff and assembled on the PI substrate via transfer printing^42–44^. Additional images, optoelectronic properties and structural details of these optical components are provided in the Methods, Figures S3, S4 and Table S1. Collectively, the LED and the detector exhibit optimal excitation and absorption characteristics that are well-suited to the optical properties of PtTFPP (Figure 1f). Figure 1g display images of a fully integrated PbtO_2_ sensing probe, which measures approximately 300 μm in width, 150 μm in thickness, and can be adjusted in length based on the depth of the targeted brain region. Upon changing the ambient gas environment from air to pure nitrogen (N_2_), the micro-probe exhibits a discernible transition from violet to red emission, attributed to the enhanced photoluminescence (PL) of PtTFPP at decreased O_2_ concentrations. Similar responses are observed when the micro-probe is inserted into a brain phantom, as shown in Figure 1h.

**Figure 1.**
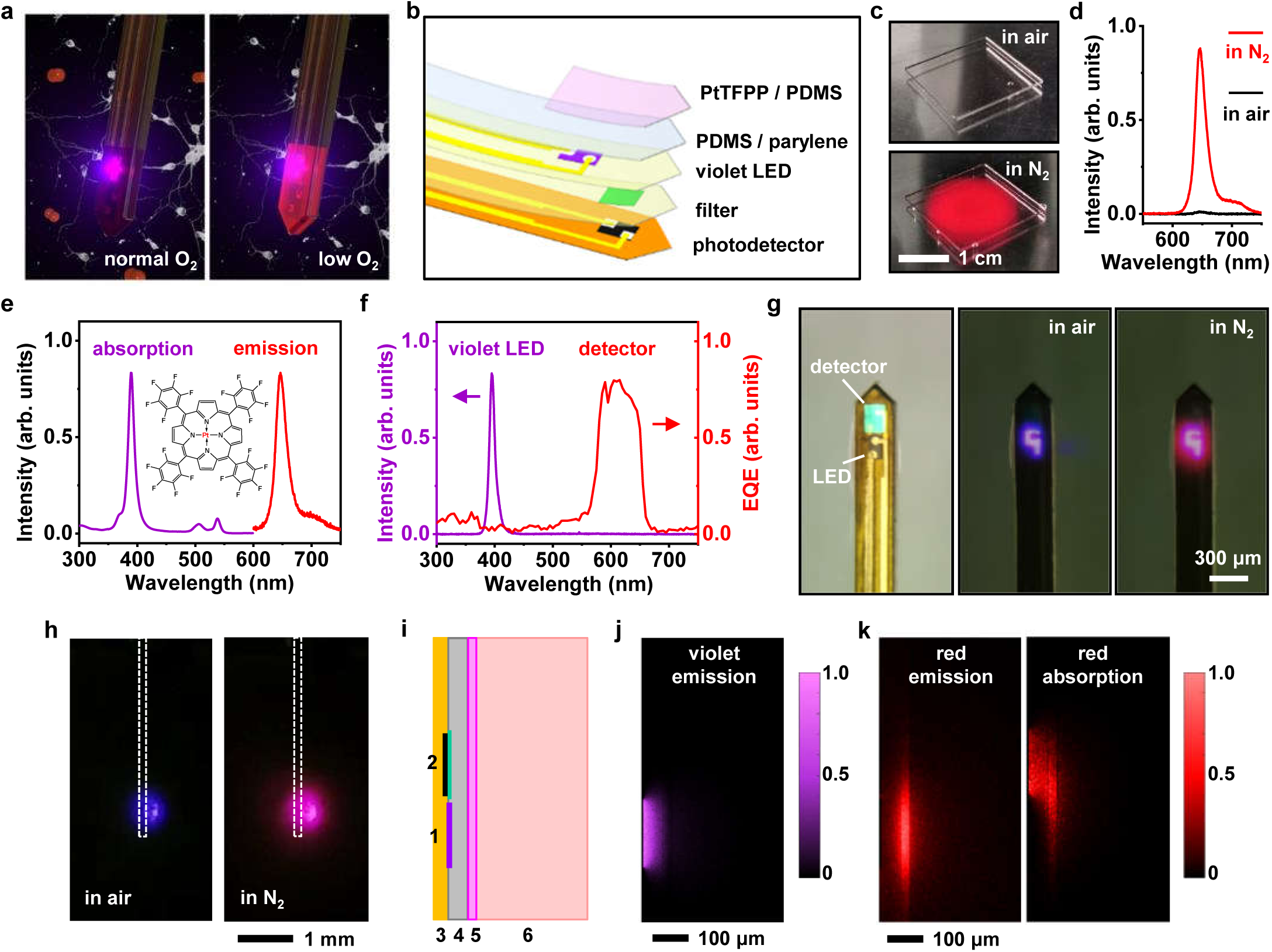
Microscale optoelectronic probe for monitoring the brain tissue oxygenation (PbtO_2_). (a) Schematic illustration of the probe implanted into the brain tissue, optically responding to the change of oxygen concentrations (left: normal condition; right: low oxygen level). (b) Enlarged view of the probe design, including (from top to bottom) a PtTFPP/PDMS sensing film, a PDMS/parylene encapsulation layer, an InGaN violet LED, a filter and an InGaP photodetector on a polyimide substrate. (c) Photographs of a PtTFPP/PDMS film excited by violet light in air (top) and in pure nitrogen (N_2_) environment (bottom). (d) PL spectra of the PtTFPP/PDMS film in air and in N_2_. (e) Absorption and PL spectra of PtTFPP in PDMS. Inset: molecular structure of PtTFPP. (f) EL spectrum of the violet LED and EQE spectrum of the detector with a filter. (g) Top-view optical images of a probe (left: LED is off; middle: LED is on in air; right: LED is on in N_2_). (h) Experimental photographs comparing light emission from a probe (side-view) implanted in a brain phantom in air (left) and in pure N_2_ (right). (i) Schematic probe structure (side-view) used in the optical modelling. (1: LED; 2: detector with filter; 3: polyimide; 4: PDMS/parylene; 5: PtTFPP/PDMS; 6: brain tissue). (j) Simulated intensity distribution of the violet excitation light from the LED. (k) Simulated red PL emission from the PtTFPP (left) and that captured by the detector (right) in N_2_.

We further establish optical models to simulate light distributions within the micro probe and surrounding tissue (Figure 1i). Figure 1j demonstrates that the violet emission from the LED is predominantly absorbed by the PtTFPP/PDMS coating and nearly no leakage of PtTFPP into the surrounding tissue. By contrast, the down-converted red phosphorene emitted by PtTFPP can penetrate more deeply into the tissue and be partially captured by the adjacent detector. Figure 1k illustrates the corresponding spatial distributions (normalized) of emitted and captured red photons. The different distributions of violet and red emissions, as well as the laterally arranged device architecture, further minimize the potential interference of violet light on the detector.

### In vitro characterization of the PbtO_2_ sensing probe

We evaluate the performance of the micro-probe in a gas chamber with varied oxygen partial pressures (pO_2_) by mixing N_2_ and O_2_ gas flows (Figure 2a). Optical signals of the LED and the detector on the micro-probe are controlled and recorded via a wireless circuit, which is described in the subsequent sections. At an LED injection current of 0.5 mA, the detector collects the PL intensity of the PtTFPP/PDMS coating at pO_2_ levels from 0 to 80 mmHg (Figure 2b). The inverse of the PL intensity is linearly dependent on pO_2_, suggesting that their relationship follows the Stern-Volmer equation (Figure 2c)^33^. The nonlinear relationship between PL intensity and pO_2_ shows that the micro-probe is more sensitive to O_2_ fluctuations at lower pO_2_ levels, which promises its potential effectiveness in monitoring cerebral hypoxia states in vivo. The measured pO_2_ detection resolutions are < 0.8 mmHg at pO_2_ = 3.8 mmHg, and < 13 mmHg at pO_2_ = 70 mmHg at room temperature (Figure S5). We also examine the micro-probe’s dynamic response to alternated O_2_ and N_2_ flows of multiple cycles (Figure 2d). When the gas flow (10 L/min) is switched between O_2_ and N_2_, recorded signal rise and decay times are 15.7 s and 0.9 s, respectively (Figure 2e). This asymmetric dynamic response is primarily ascribed to the nonlinear relationship between PL intensity and pO_2_ (Figure 2b). The transient response of our micro-probe system is faster than previously reported techniques based on optical sensors and Clark electrodes^28, 39, 40, 45, 46^. Additionally, the PL quenching mechanism in PtTFPP ensures that the micro-probe possesses high O_2_ selectivity over other gases including N_2_ and CO_2_ (Figure 2f). After the pretreatment, the micro-probe can continuously monitor pO_2_ for over 2 h without any significant photobleaching (Figure 2g). Besides photostability, the micro-probe also exhibits desirable thermal stability, with a change of only ∼0.85% in PL signal readout per degree Celsius from 28 °C to 38 °C (Figure 2h). During in vitro and in vivo operation, the LED is driven by a pulsed current of 0.5 mA, which only causes a temperature rise of less than 1 °C (Figure S6). Therefore, the device’s heating generates minimal effects on measurement accuracy and normal tissue function. Finally, the absorption of the PtTFPP/PDMS coating remains unchanged when immersed in phosphate-buffered saline (PBS) solution for over a month (Figure 2i and Figure S7), which indicates the absence of any significant leakage of phosphors and promises chronic operation stability.

**Figure 2.**
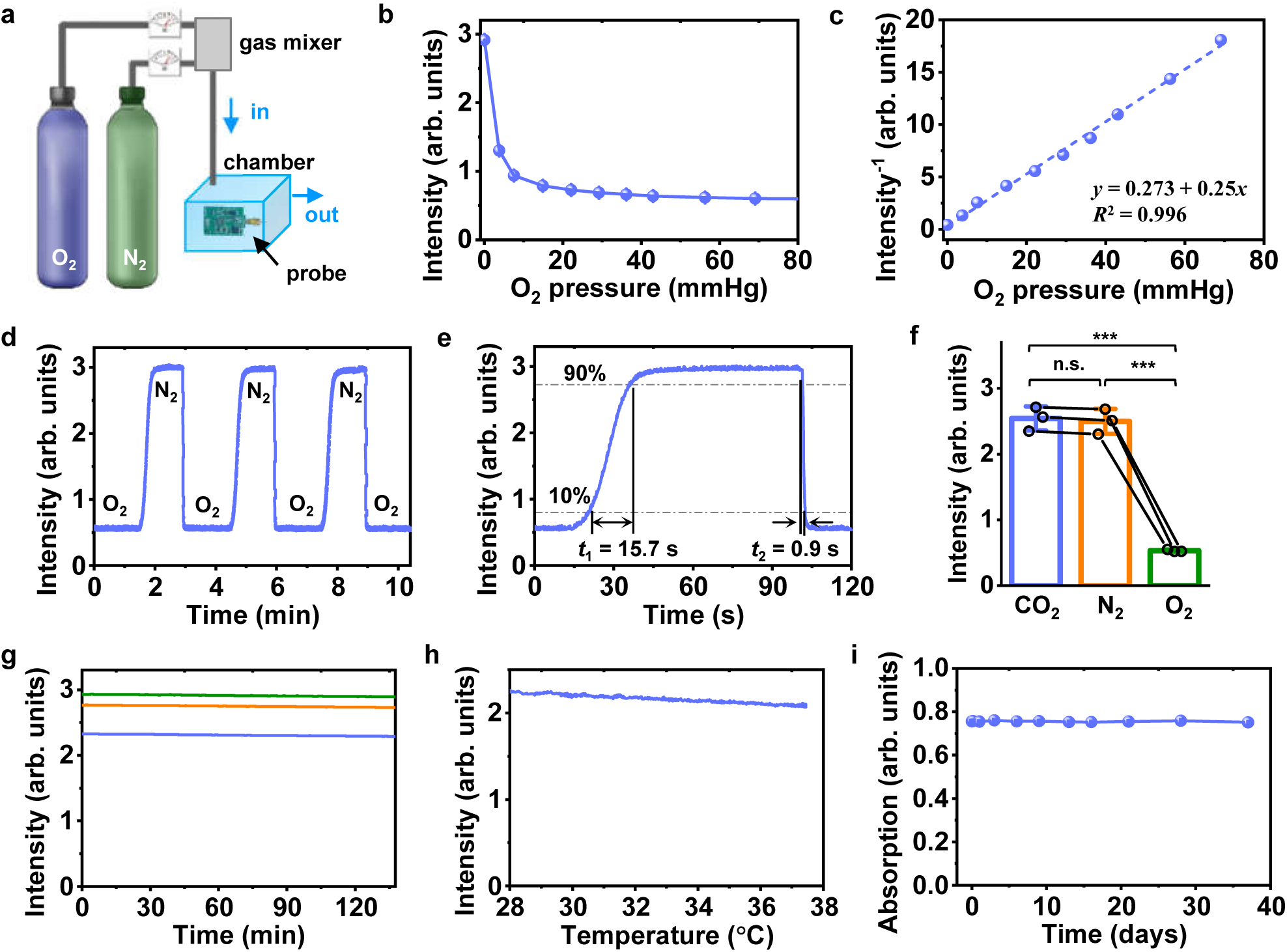
In vitro characterization of the PbtO_2_ sensing probe. (a) Illustration of the experiment setup for measuring the probe in response to different O_2_ levels. The gas chamber has a size of 10 cm × 10 cm × 6 cm, and gas flows of N_2_ and O_2_ are varied. (b) Recorded PL intensities at different O_2_ partial pressures. (c) Calibration curve showing the linear relationship between the inverse of PL intensities and O_2_ partial pressures. (d) Dynamic response of the probe at alternated O_2_ partial pressures (from 0 to 760 mmHg). (e) Partially enlarged curve of (d), indicating a rise time *t*_1_ = 15.7 s and a decay time *t*_2_ = 0.9 s. (f) Recorded signals of the probe responding to different gases (CO_2_, N_2_ and O_2_, pressure ∼1 atm or ∼760 mmHg). One-way ANOVA and LSD post-hoc comparison are performed (*n* = 3 probes, *** *P* < 0.001, n.s. *P* > 0.05). (g) Signal stability of three different probes (LED injection current = 0.5 mA), measured at room temperature in pure N_2_. (h) The probe response at different temperature (from 28.5 °C to 37.5 °C). (i) Chronic stability of a PtTFPP/PDMS film. The film is immersed in PBS for more than a month and the peak absorption (at 390 nm) is measured at different days.

### Real-time, in vivo PbtO_2_ monitoring with the wireless probe

For continuous PbtO_2_ monitoring in living animals, we implement the implantable optoelectronic probe with a customized wireless control circuit. Shown in Figure 3a, key components in the circuit involve an LED driver, an operational amplifier (AMP), a Bluetooth system on chip (SoC), and a rechargeable battery. An external computer wirelessly controls the devices and collects optical signals via Bluetooth communication. Data are converted to corresponding PbtO_2_ levels based on in vitro calibrations (Figure 2c). The circuit module has a compact size of 12.3 × 17.3 mm^2^ and a weight of 1.8 g (Figure 3b) and is head-mounted on a freely-moving mouse for continuous monitoring of PbtO_2_ (Figure 3c). More details about the battery-powered circuit are presented in Figures S8, S9, and Table S2. To assess the sensing capability, we conduct acute tests on a mouse by dynamically varying the fractions of inspired oxygen (FiO_2_) using a mixture of inhaled N_2_ and O_2_ (Figure 3d). Our wireless micro probe and a commercial fiber-optic sensor (NeoFox-GT, Ocean Insight, FOSPOR AL300 probe) are inserted into the same brain region (bilateral hippocampus, at a depth of ∼2.2 mm) to simultaneously record PbtO_2_ signals. The data collected by our wireless micro-probe agree well with those collected by the commercial fiber-optic sensor (Figure 3e), suggesting these two systems exhibit comparable performance. It should be noted that a FiO_2_ of 7.4% corresponds to a recorded PbtO_2_ level of less than 10 mmHg, which indicates a severe cerebral hypoxia event^47^. Furthermore, we employ the micro-probe to record real-time PbtO_2_ in mice during acute ischemia/reperfusion induced by clamping and declamping their bilateral carotids (Figure 3f). Recorded PbtO_2_ levels decrease to less than 3 mmHg after clamping for about 1 min, and recover after reperfusion (Figure 3g). Another validation test monitors and compares PbtO_2_ levels for mice in awake and anesthetic states. The micro-probe records significantly higher PbtO_2_ levels in mice under isoflurane anesthesia. Isoflurane, a commonly used inhaled anesthetic, is also known to be a potent vasodilator^48^, and our findings indicate that brain tissue oxygenation is significantly enhanced under isoflurane (2%) anesthesia, consistent with previous studies^49^. Finally, and most importantly, the wireless circuit modules support the acquisition and transportation of data from multiple micro-probes in multiple animals, which is a feature hardly realized with the tethered fiber-optic system. As shown in Figure 3i and Movie S1, the system simultaneously monitors PbtO_2_ levels of three behaving mice in the same chamber without affecting their locomotion. When supplied with varied FiO_2_, the three animals exhibit similar PbtO_2_ fluctuations (Figure 3j). These explorations demonstrate distinct advantages of our wireless micro-probe in studying real-time PbtO_2_ activities associated with complex animal behaviors and interactions.

**Figure 3.**
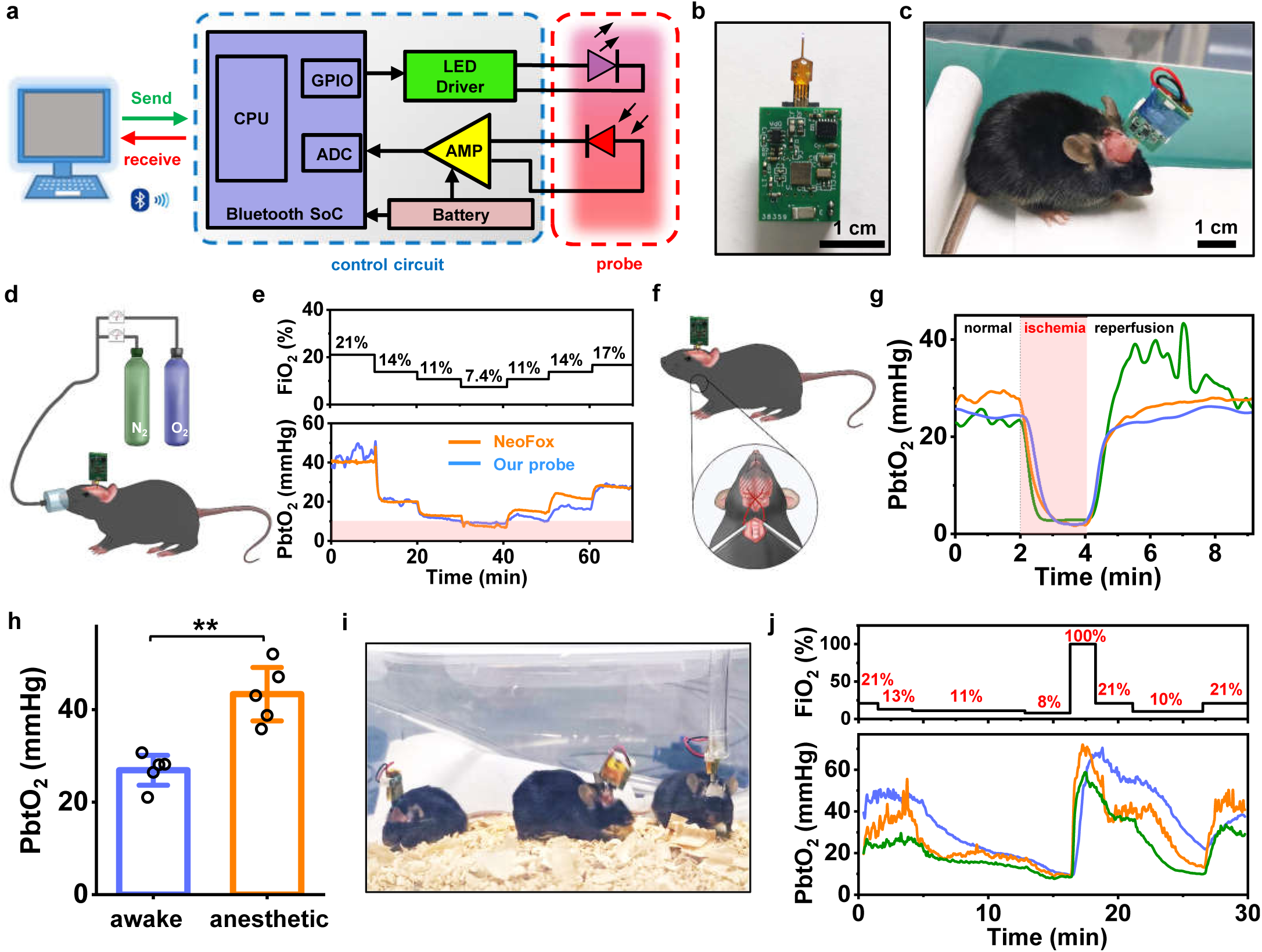
The probe combined with a wireless control circuit continuously monitors PbtO_2_ in the brain of behaving mice. (a) Block diagram of the wireless system, comprised of a computer, a battery-powered control circuit and an implantable probe. (b) Image of a probe connected with a wireless circuit. (c) Photo showing a behaving mouse with an implanted probe and the head mounted circuit. (d) Scheme of the setup showing that a wireless probe monitors PbtO_2_ of a mouse provided with different fraction of inspired oxygen (FiO_2_). (e) Dynamic change of FiO_2_ (top) and PbtO_2_ results (bottom) simultaneously detected by our wireless probe and a commercial fiber recording system (Ocean Optics NeoFox). (f) Scheme showing a probe monitors PbtO_2_ of a mouse during acute ischemia/reperfusion induced by clamping and declamping its bilateral carotid. (g) Recorded PbtO_2_ results during acute ischemia/reperfusion (*n* = 3 mice). (h) Comparison of PbtO_2_ results measured in mice during their awake and anesthetized conditions. A paired t-test is performed (*n* = 5 mice, ** *P* < 0.01). (i) Photo of three freely moving mice with their PbtO_2_ simultaneously monitored by wireless probes. (j) Dynamic change of FiO_2_ (top) and PbtO_2_ results (bottom) simultaneously detected in the three mice.

### The wireless probe records PbtO_2_ in the mouse seizure model induced by electrical stimulations

In the mammalian brain, the PbtO_2_ level strongly correlates with local neural activities. Intense neural activities usually lead to an increase in local metabolism and oxygen consumption, resulting in decreased PbtO_2_ levels^17, 50^. Conversely, abnormalities in PbtO_2_ levels cause dysfunction of neural activities and impair brain function^51, 52^. Utilizing our micro-probe, we investigate local PbtO_2_ variations in response to electrical stimulations in the deep brain. Schematically shown in Figure 4a, stimulating and recording electrodes insert into the hippocampus (CA1) of mice. Meanwhile, the optoelectronic micro-probe is implanted into the ipsilateral hippocampus (CA3) to continuously capture PbtO_2_ changes. The surgical procedures are described in detail in Figure S10, and the coronal section stained with Nissl shows the lesion region created by the micro-probe (Figure S11). The hippocampus is a brain region with high metabolic activity and specialized functions related to memory and learning behaviors, making it particularly sensitive to changes in PbtO_2_ levels^53^. The electrical stimulation (kindling) protocol involves a biphasic pulse train with a current of 0.4 mA, a pulse frequency of 60 Hz, and a pulse width of 1 ms, administered for a duration of 1 s. Figure 4b and Movie S2 display an instance of a dramatic seizure event in a mouse, whose motion behaviors, local field potentials (LFPs) in CA1 and PbtO_2_ levels in CA3 are dynamically monitored simultaneously. In this example, a 1-s pulsed kindling in CA1 elicits prominent afterdischarges (ADs), appearing in the form of intensified LFP oscillations. The ADs sometimes accompany severe convulsion in the animal. The LFP bursts last for about 40 s, and then the micro-probe records dramatically decreased PbtO_2_ at the end of the animal convulsion, suggesting that intense neural activities cause increased local oxygen consumption. It is also noted that not every kindling can trigger AD and associated PbtO_2_ drops (Figure S12). Statistically, the CA3 region experiences severe hypoxia (PbtO_2_ < 10 mmHg) approximately 30 s after each AD, and recovers within about 60 s (Figure 4c). This temporal relationship suggests that the kindling induces excessive neural activities locally, elicits convulsive behaviors, and eventually leads to more oxygen consumption by the neurons^54^. The resulting hypoxia then suppresses neural activity and terminates the convulsive behavior in mice^55, 56^.

**Figure 4.**
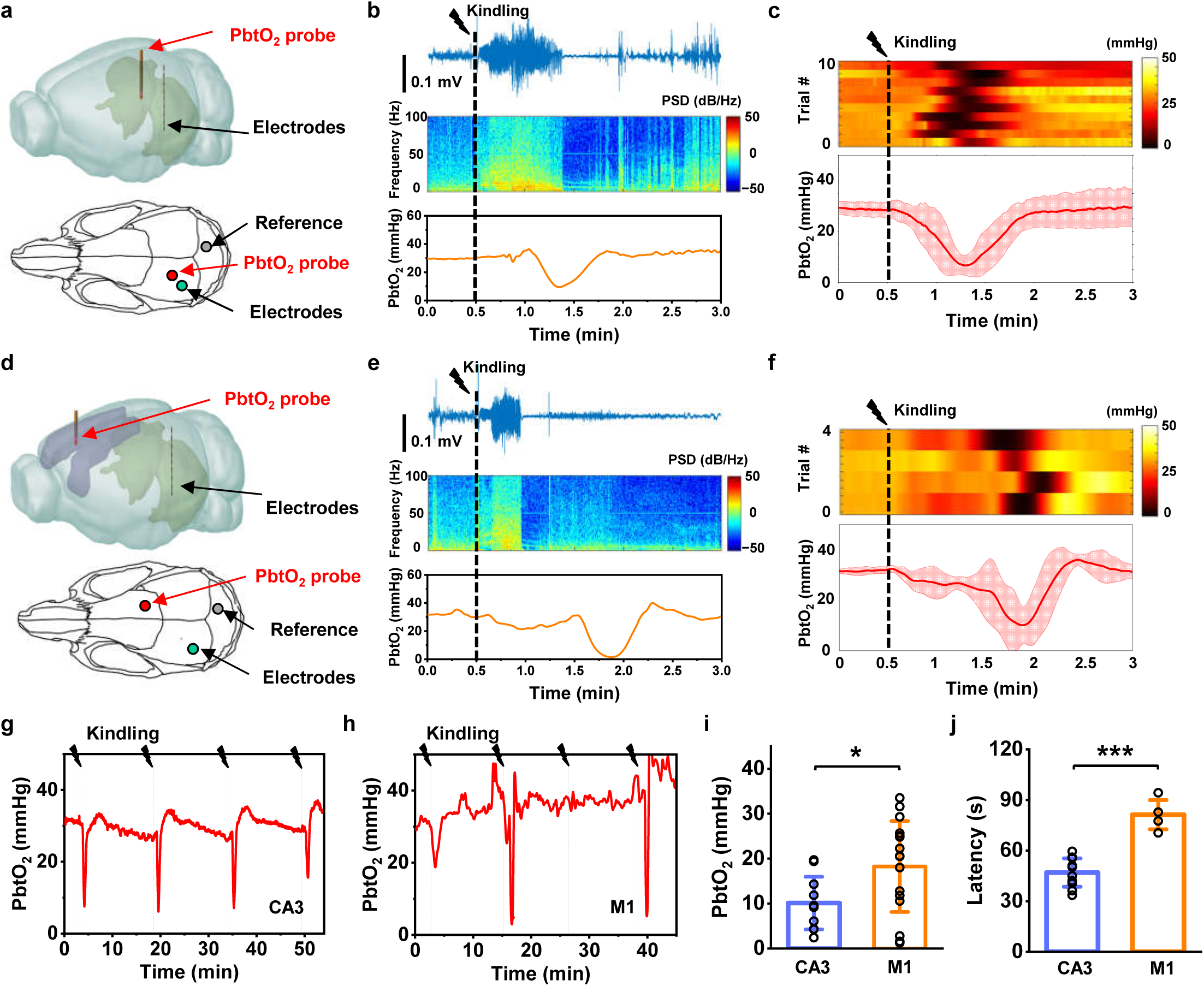
The wireless probe records PbtO_2_ changes in different brain regions of mice under electrical stimulations. (a) 3D (upper) and 2D (lower) schematic models showing positions of the PbtO_2_ sensing probe and the stimulating/recording electrodes implanted in the ipsilateral hippocampus of mice (CA3 and CA1, respectively). (b) Simultaneously recorded electrophysiological activities in CA1 and PbtO_2_ in CA3 under kindling from a mouse. Top: LFP trace. Middle: power spectral density (PSD) of LFP. Bottom: PbtO_2_. (c) Dynamic response of PbtO_2_ recorded in CA3 immediately after kindling. Top: Heatmap of 10 individual trials from *n* = 10 mice. Bottom: Average PbtO_2_ signals. The solid lines and shaded areas indicate the mean and s.e.m., respectively. (d) 3D (upper) and 2D (lower) schematic models showing positions of the PbtO_2_ sensing probe and the stimulating/recording electrodes implanted in the contralateral primary motor cortex (M1) and the hippocampus (CA1) of mice, respectively. (e) Simultaneously recorded electrophysiological activities in CA1 and PbtO_2_ in M1 under kindling from a mouse. Top: LFP trace. Middle: power spectral density (PSD) of LFP. Bottom: PbtO_2_. (f) Dynamic response of PbtO_2_ recorded in M1 immediately after kindling. Top: Heatmap of 4 individual trials from *n* = 3 mice. Bottom: Average PbtO_2_ signals. The solid lines and shaded areas indicate the mean and s.e.m., respectively. (g) Continuously monitored PbtO_2_ in CA3 of a mouse during sequential electrical stimulation (kindling) for multiple times. (h) Continuously monitored PbtO_2_ in M1 of a mouse during sequential electrical stimulation (kindling) for multiple times. (i) Comparison of the lowest PbtO_2_ values recorded in ipsilateral CA3 (10 trials, *n* = 5 mice) and contralateral M1 (16 trials, *n* = 4 mice) after kindlings in CA1. Statistical analysis is based on independent samples *t*-test. * *P* < 0.05. (j) Comparison of the latency time from kindlings in CA1 to reaching the lowest PbtO_2_ in CA3 (10 trials, *n* = 5 mice) and M1 (4 trials, *n* = 3 mice). Statistical analysis is based on independent samples *t*-test. * *P* < 0.05, *** *P* < 0.001.

Furthermore, we utilize the micro-probe to record PbtO_2_ in a distant region, the primary motor cortex (M1), during hippocampal kindlings. In this experiment, the micro-probe is in the contralateral M1 while the stimulating/recording electrodes are still placed into CA1 (Figure 4d). As shown in Figure 4e, Figure 4f and Movie S3, the kindling-induced ADs can also elicit hypoxia in M1. While almost every kindling in CA1 can induce a corresponding hypoxia event in the ipsilateral CA3 (Figure 4g), similar PbtO_2_ drops are not always observable in M1, and they occur in approximately 25% of the trials (16 trials from 4 mice) (Figure 4h). Compared with the results in CA3, similar hypoxia events observed in M1 are less pronounced (Figure 4i) and have a higher latency (Figure 4j). The high latency in M1 may be attributed to the longer electrical transfer time from CA1 to M1, while the lower probability can be attributed to the fact that the transfer pathway is not exclusive.

The above PbtO_2_ monitoring results are different from those data collected by fMRI, which is typically more sensitive to increased blood flow and hemodynamic response in the seizure occurring area^57, 58^. Such a divergence clearly reveals that blood oxygenation and PbtO_2_ are two distinct metabolic indicators. While the former concentrates more on the oxygen supply within the blood, the latter manifests the oxygen consumption of local cells and tissue.

### A fully implanted, battery-free probe for PbtO_2_ monitoring in rats

The above-mentioned experiments clearly demonstrate that the wireless micro-probe has the capability of real-time PbtO_2_ detection and associates the results with certain cerebral abnormalities. A further technical advance involves the development of a wireless, battery-free circuit module that can be fully implanted subcutaneously, as detailed presented in Figures 5a, S13, and S14. Compared with the battery-powered probe, such a battery-free system offers multiple advantages^59, 60^: (1) All the electrical components mount on a thin-film, PI-based printed circuit board (thickness 130 μm) instead of a rigid one, improving its flexibility; (2) An inductive coil replaces the battery for power harvesting, reducing the system’s weight by ∼1 g; (3) The inductively coupled power transfer strategy enables longer-lasting data collection without interruptions due to battery replacements. Protected by water-proof encapsulant, the entire circuit can be fully implanted under the scalp of behaving rats (Figures 5b, 5c, and S15). X-ray computed tomographic (CT) images reveal the location of the PbtO_2_ probe within the rat brain as well as the position of the circuit module (Figure 5d). As the power transverter, a looped antenna receives currents from an external radio frequency (RF) source and generates uniform magnetic fields in an animal behaving enclosure (Figures 5e and S16). Under normal operation, the circuit experiences a temperature rise of less than 3.5 °C, ensuring its safety within the animal body (Figure S17). The proof-of-concept study presented in Figure 5f demonstrates the wireless and continuous PbtO_2_ monitoring in a rat in response to FiO_2_ changes for over 50 mins. The recorded PbtO_2_ variations with the battery-free micro-probe are similar to those responses collected in mice with the battery-powered system.

**Figure 5.**
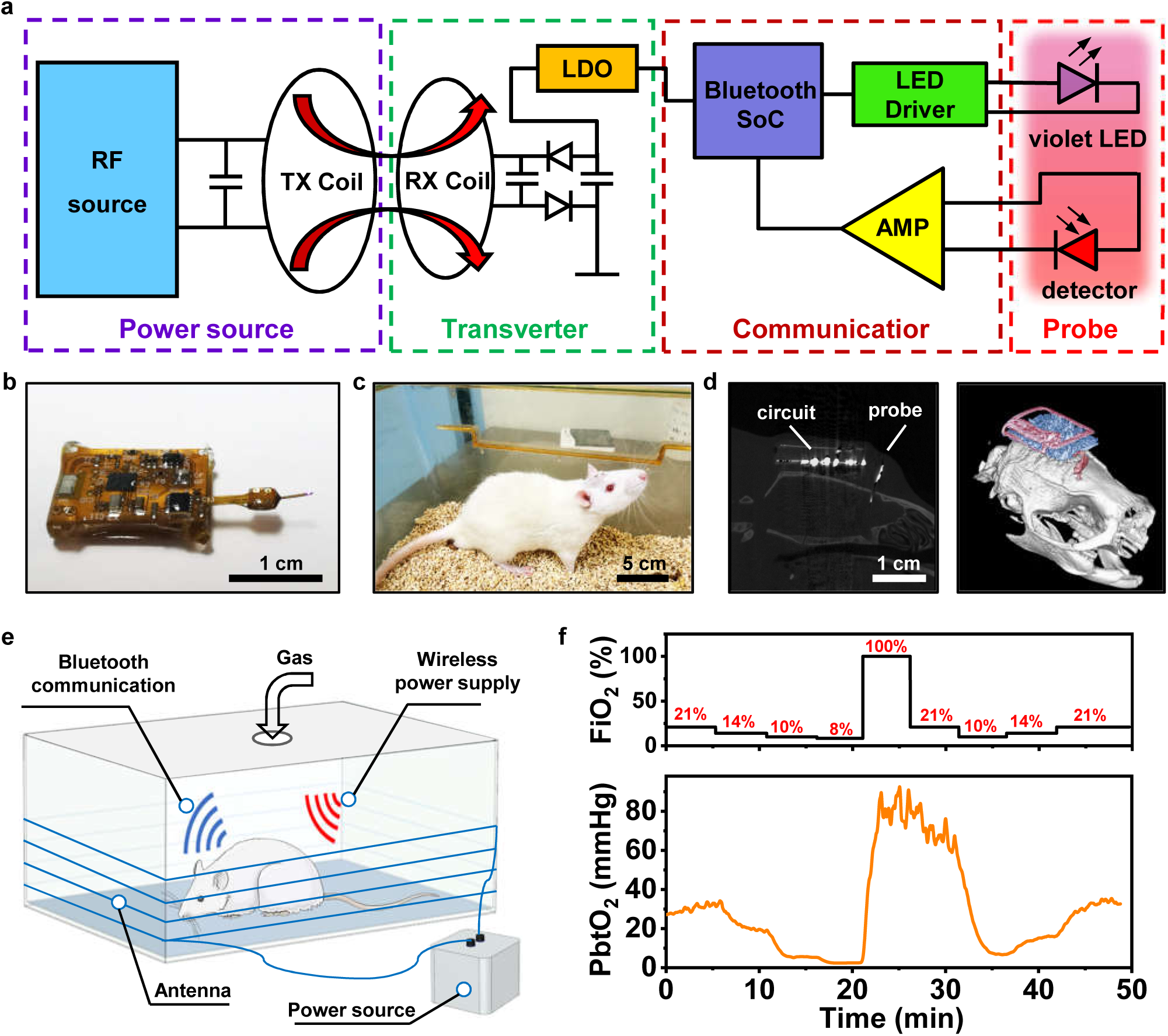
A fully implanted, battery-free probe system monitors PbtO_2_ in the brain of living rats. (a) Block diagram of the system, including an RF power source, a receiving coil, a Bluetooth communication module and an implantable probe. (b) Image of a probe connected with the battery free circuit. (c) Photo showing a freely moving rat with a fully implanted probe system. (d) Sagittal CT image indicating the location of the implanted system in a rat (left), and reconstructed 3D CT image showing the position of the device relative to the skull (right). (e) Schematic illustration of the setup for monitoring PbtO_2_ in rat. (f) Dynamic change of FiO_2_ (top) and PbtO_2_ results (bottom) monitored in a rat.

## DISCUSSION

To summarize, we establish a wireless optoelectronic probe system, which can be powered by either a battery or inductively coupled coils, to continuously monitor PbtO_2_ in a living animal brain. Table S3 compares characteristics of our micro-probe with those in previous reports. Different from those probe systems that sense blood oxygen^21, 61, 62^, our micro-probe explores tissue oxygenation through PbtO_2_, which has more direct relevance to local neural activities. The utilization of microscale devices guarantees a small footprint and minimize the lesion area in the brain tissue. The thin film PtTFPP/PDMS coating also reduces the response time to less than 1 s. Experiments in a rodent model manifest the relationship among electrical stimulations, PbtO_2_ levels and epileptic activities. During the seizure occurrence, the observed PbtO_2_ changes are clearly distinct from previously recorded blood oxygenation signals by fMRI, which is performed accompanied by optogenetic stimulation^58^. In this study, we monitor PbtO_2_ signals in a seizure model induced by electrical stimuli. In the future endeavor, it will be more beneficial to investigate PbtO_2_ before, during and after spontaneous seizure occurrences. While here our micro-probe captures the decreased PbtO_2_ level after ADs and convulsive events (Figure 4), previous works show that hypoxia can elicit seizure as well ^17, 63^, which implicates that monitoring PbtO_2_ levels using the micro-probe could also potentially serve as an early indicator for the development of severe epilepsy. Besides PL intensity recording, advanced circuit systems can enable PbtO_2_ detection based on PL lifetime changes, which are less susceptible to photobleaching of dyes and potentially provide more accurate measurements^33^. Meanwhile, the combination of electrophysiological activities, PbtO_2_ and blood oxygenations will foster a more profound understanding of the neurovascular and neurometabolic coupling^64, 65^. In the future, the PbtO_2_ probe can integrate with other microscale devices for detecting electrophysiological signals^66^, calcium fluorescence^67, 68^, neurotransmitters^69, 70^ and temperature^71, 72^, to realize multi-modal neural sensing. Additionally, the PbtO_2_ sensing technique can collaborate with optogenetic and non-genetic stimulations^73, 74^, to achieve closed-loop interrogation. The wireless circuits facilitate studies among multiple untethered targets under complex behaviors, which are unavailable with wired devices. In practice, both battery-powered and battery-free circuit schemes can be feasible for clinical applications. The presented technique clearly implicates its potential utility in neuroscience research and medical diagnostics.

### Methods

#### Device Fabrication

Details about device structures and fabrication processes are provided in the supplementary information, Figs. S1–S4 and Tables S1. The InGaP photodetector structure is grown on a GaAs substrate, and the InGaN violet LED structure is grown on sapphire, both by metal-organic chemical vapor deposition (MOCVD). Both of the detector and the LED are fabricated via a standard photolithographic process, and freestanding, thin-film microscale devices are formed by selectively etching the Al_0.95_Ga_0.05_As sacrificial layer (for detector)^67^ or ultraviolet laser liftoff (for LED)^75^. The multilayered TiO_2_/SiO_2_ longpass filter is deposited on GaAs substrate by ion beam–assisted sputtering, with freestanding, thin-film filters formed by femtosecond laser milling and wet etching^42^. A detector, an optical filter and a violet LED are sequentially transferred onto the flexible substrate with PDMS stamps^76^, followed by insulation with SU-8 and metallization. Laser milling (ultraviolet laser 365 nm) defines the needle shape (width ∼300 μm, length ∼3 mm). Sputtered Cr/Au/Cu/Au (6 nm/100 nm/600 nm/150 nm) layers serve as interconnect contacts for the devices. Laser milling is used to form the shape of micro-probes. After encapsulated by PDMS and parylene C, an oxygen sensing film (PtTFPP/PDMS) is dip coated on the tip of the probe.

#### Device Characterization and Modeling

##### Characterization of the PbtO_2_ sensing probe

In vitro characterizations of the PbtO_2_ monitoring probe are performed in a gas chamber with a size of 10 cm × 10 cm × 6 cm, flowed from a gas mixer with two gas flowmeters connected to pure N_2_ (range: 0∼25 L/min, resolution: 2.5 L/min) and pure O_2_ (range: 0∼1 L/min, resolution: 0.05 L/min). The pO_2_ level is controlled by the two gas flowmeters. Prior to calibration, the probe undergoes a photobleaching pre-treatment in the pure N_2_ environment, by supplying the LED with a constant current of 0.5 mA for 30 min. During in vitro test, the probe is operated with a sampling frequency of 4 Hz and a driver current of 0.5 mA at a pulse width of 10 ms. The thermal stability test is performed in a PBS solution containing Na_2_SO_3_ (5% w/v).

##### Optoelectronic characterizations

The emission spectra of the oxygen-sensitive film and the LED are measured with a high-resolution spectrometer (HR2000+, Ocean Optics). The absorption spectra of the oxygen-sensitive film and transmission spectra of the filter and the polyimide film are measured using a spectrophotometer (Cary5000, Agilent Technologies). Current– voltage characteristics of the LED and photodetector are recorded by a Keithley 2400 source meter. External quantum efficiencies of the LED and the detector are determined using a spectroradiometer with an integrating sphere (LabSphere Inc.). For in vitro tests, micro-probes are implanted into a brain phantom made of agarose (0.5% w/v), hemoglobin (0.2% w/v), and intralipid (1% w/v) mixed in PBS (98.3% w/v). After stirring, the mixed liquid is heated to boiling, then cooled to room temperature and form a gel.

##### Optical modeling

A ray-tracing method based on Monte Carlo simulations (TracePro free trial version) simulates the light distribution in the brain. In the model, an LED with a size of 190 μm × 110 μm × 10 μm emits light at a wavelength of 395 nm and a detector (185 μm × 125 μm × 3 μm) is placed adjacent to the LED. The LED emission has a Lambertian distribution and is modeled with 10^6^ rays. A layer of polymer with a thickness of 50 μm and a layer of oxygen-sensitive film with a thickness of 25 μm are coated on both devices. The refractive indices of the polymer and oxygen-sensitive film are 1.44 at 395 nm and 1.41 at 650 nm. Fluorescence property is applied on the oxygen-sensitive film with a conversion efficiency of 0.19, a peak molar extinction of 300000 M^-1^cm^-1^ and a molar concentration of 0.85 mM. The probe is inserted into the brain tissue (size 5 × 5 × 5 mm^3^) with absorption coefficients of 0.75 /mm and 0.08 /mm, and scattering coefficients of 30 /mm and 33 /mm, for violet (395 nm) and red (650 nm) light, respectively. The tissue has an anisotropy factor of 0.85 and a refractive index of 1.36.

##### Thermal measurement and modeling

The temperature distributions on the surface of the probe and the circuit board are mapped with an infrared camera (FOTRIC 228). The emissivity values of the probe surface, the surface of the battery-free circuit and the surface of the battery-powered circuit are set to 0.95, 0.95 and 0.80. A 3D steady-state heat transfer model is established by finite element analysis (Comsol Multiphysics). Materials and corresponding parameters (density, thermal conductivity and heat capacity) used in the model are as follows: parylene (1.11 g/cm^3^, 0.086 W/m/K, 3.5 J/g/K), PDMS (0.965 g/cm^3^, 0.15 W/m/K, 1.46 J/g/K), polyimide (1.4 g/cm^3^, 0.15 W/m/K, 1.1 J/g/K), Cu (8.96 g/cm^3^, 400 W/m/K, 0.385 J/g/K), SU-8 (1.2 g/cm^3^, 0.2 W/m/K, 1.5 J/g/K). The boundary condition is natural heat convection to air at 25 °C. The LED serves as the heat source, with the input thermal power estimated by *P* = *V* × *I* × (1 − EQE), where *V*, *I,* and EQE are the measured voltage, current, and corresponding external quantum efficiencies for LEDs. The measured and simulated thermal distribution of the probe surface is shown in Figure S6.

#### Wireless Circuit Design

A customized circuit module is designed to wirelessly operate the LED and receive signals from the detector, and a computer or a mobile phone controls the monitoring system and receives PbtO_2_ signals. The schematic diagram and printed circuit board layout of the battery-powered circuit design are depicted in Figure S8 and Figure S9, respectively. A programmable microcontroller (nRF52832, Nordic Semiconductor) operates at 2.4 GHz for wireless data communication. An LED driver (LTC3212, Analog Device) with an external resistor supplies a stable current to the LED. The photocurrent of the detector is amplified using an operational amplifier (OPA391, Texas Instruments), which is then digitized by the microcontroller’s analog-to-digital converter for wireless transmission. A linear regulator (NCP161, ON Semiconductor) with a fixed internal output voltage (3.3 V) ensures a stable power supply. The battery powered circuit has a size of 17.3 × 12.3 mm^2^ and a weight of 1.8 g (including a rechargeable lithium-ion battery with a weight of 1.2 g). The battery has a capacity of 45 mAh and can continuously work for more than 8 hours. The circuit is connected to the implanted probe via a flexible connector.

The design of the battery-free circuit is similar to that of the battery-powered circuit, except that it is powered through inductive coupling. The schematic diagram and layout of the fully implanted battery-free circuit are displayed in Figures S13 and Figure S14, respectively. The flexible printed circuit board is based on PI/Cu foils, and an inductive coil with 7 turns is used for the power supply. Two capacitors in parallel (∼62 pF) are employed to tune the power harvesting antenna. The half-bridge rectifier is made up of two Schottky diodes (Skyworks) and a capacitor (2.2 µF). Located in a cage enclosed by a copper wire antenna with four loops, the PbtO_2_ of behaving rats can be continuously monitored with the fully implanted micro-probe. Electromagnetic finite element analysis is applied to simulate the magnetic field distribution inside a 23 × 16 × 6.5 cm^3^ enclosure (Figure S16). The antenna operates at 13.56 MHz, and the output power is 4 W. The maximum temperature rise of the wireless circuit tested at room temperature is less than 3.5 °C (Figure S17).

#### Biological Studies

##### Animals

All animal procedures are approved by the Institutional Animal Care and Use Committee (IACUC) at Tsinghua University. Adult (8–12 weeks) male wild-type C57BL/6N mice and adult male Sprague–Dawley (SD) rats (350g–550g) purchased from the Vital River Laboratory (Animal Technology, Beijing, China) are used and housed in groups (3–5 mice per cage, 1–2 rats per cage) under standard conditions. Animals are maintained on a 12 h light/dark cycle at 22–25 °C and provided with access to food and water ad libitum.

##### Stereotaxic Surgery

Adult wild-type mice are anesthetized with an intraperitoneal injection of Avertin (280 mg/kg) before surgery and then placed in a standard stereotaxic instrument for surgical implantation (Figure S10). After shaving the scalp and exposing the skull, small holes with a diameter of ∼0.5 mm are drilled with a micromotor drill on the skull. For mice provided with electrical stimulations, PbtO_2_ sensing probes are slowly implanted in the hippocampal CA3 (AP: −1.8 mm, ML: ±1.86 mm, DV: −2.25 mm) or M1 (AP: 1.15 mm, ML: 1.7 mm, DV: −2 mm), and a small amount of dental cement fixes the probe. The stimulating/recording electrodes are constructed of twisted Teflon-coated tungsten wires (Cat No. 796000, A-M Systems) and targeted to the hippocampal CA1 region (AP: −3.0 mm, ML: −2.9 mm, DV: −3.2 mm). The stainless steel screw works as the reference electrode and implants on the skull. Finally, more dental cement is applied to fix all devices on the skull (Figure S10).

The battery-free circuit is fully implanted in SD rats after anesthetized with an intraperitoneal injection of mixed Zoletil and Xylazine (8 and 16 mg per kg). After placed in a standard stereotaxic instrument for surgical implantation, an incision of the scalp is created to expose the skull. Then, 0.3% hydrogen peroxide is applied to clean the skull, and a small hole with a diameter of ∼0.5 mm was drilled. The probe is implanted into the brain (AP: 3.6 mm, ML: 2.0 mm, DV: −2.9 mm), and dental cement is applied to fix the device on the skull followed by suturing the scalp (Figure S15).

In vivo measurements on mice and rats are conducted at least 4 days after surgery to allow animals to recover.

##### In vivo Pbt*O_2_ monitoring*

Clamping and declamping mice bilateral carotid are performed on the mice anaesthetized with Avertin (280 mg/kg). Under anesthesia, the wireless circuit connects to the probe fixed on the mouse skull, then the mouse is placed in a supine position, and a small incision is made in the middle of the anterior neck. Next, the muscles under the skin are removed, and the bilateral common carotid artery (CCA) is exposed. The left CCA is ligated by a small artery clip, and then the right CCA is ligated. After monitoring for about 2 mins, the clips are removed to restore the blood flow.

Micro-probes monitor the PbtO_2_ of behaving mice in varied FiO_2_. One or multiple mice are placed in a gas chamber with a size of 16 cm × 23 cm × 13.5 cm, and the inflowing gas is controlled by two flowmeters connected to pure N_2_ and pure O_2_. The ratio of N_2_ and O_2_ flow rates are varied to achieve different pO_2_.

PbtO_2_ of awake mice is measured 5 min after they are placed a gas chamber with an airflow of 2 L/min, and PbtO_2_ of their anesthetized states is measured 5 min after flowing air mixed with 2% isoflurane.

Electrical stimulations/kindlings are applied on mice in the hippocampal CA1 region at least 4 days after surgery. Sequential biphasic kindlings (1-ms pulse width, 60 Hz for 1 s) are applied with a time interval of no less than 10 min for 8 times per day until seizure occurs (usually 2∼4 days after kindlings). LFP recording and electrical stimulations are accomplished using an electrophysiological recording system (alphaRS Pro, Alpha Omega Engineering), and the implanted micro-probe wirelessly records the PbtO_2_ signals simultaneously.

##### Nissl staining

Mice are anesthetized with Avertin (280 mg/kg) and perfused intracardially with PBS and 4% paraformaldehyde. The brains are post fixed in 4% paraformaldehyde overnight. Coronal brain paraffin sections are taken with a thickness of 5 µm, and then deparaffinized with xylene twice (for 20 minutes each). These sections are rinsed with anhydrous alcohol twice (for 5 minutes each) and 75% alcohol for 5 minutes. Then, they are rinsed with tap water and distilled water. After immersed in 0.03% toluidine blue solution (pH = 1.5) for 2–5 minutes, samples are washed with tap water. The sections are cleared in xylene for 10 min and mounted with permanent mounting media.

## Acknowledgements

This work is supported by Beijing Municipal Natural Science Foundation (Z220015), National Natural Science Foundation of China (NSFC) (52272277, 62005016), Center for Flexible Electronics Technology at Tsinghua University.

## Author contributions

X. C., D. S., Z. L., W. Z., Y. X., X. F., and B. Z. performed device design, fabrication and characterization. X. C., H. Z., Z. Y., Z. X., and H. D. designed and tested circuits. X. C., P. W., Q. L., D. S., Y. D., C. W., and Y. W. designed and performed biological experiments. P. W., C. W., L. Y., H. P., H. D., G. Z., and X. S. provided tools and supervised the research. X. C., H. D., and X. S. wrote the paper in consultation with other authors.

## Data and materials availability

All data needed to evaluate the conclusions in the paper are present in the paper and/or the Supplementary information.

## Supplementary Information

### Preparation of individual components in the micro-probe

The layout of the microscale oxygen sensing probe includes (from top to bottom) a platinum(II)-5,10,15,20-tetrakis-(2,3,4,5,6-pentafluorophenyl)-porphyrin / poly(dimethylsiloxane) (PtTFPP/PDMS) sensing film (thickness: ∼25 μm), a PDMS/parylene C encapsulation layer (thickness: ∼50 μm), an SU-8 layer (thickness: 5 μm), an indium gallium nitride (InGaN) violet LED (size: 190 μm × 110 μm × 10 μm), a polyimide (PI) layer (thickness: 8 μm), an optical filter layer based on multilayer titanium dioxide (TiO_2_) and silicon dioxide (SiO_2_) (size: 220 μm × 150 μm × 6 μm), a PI layer (thickness: ∼4 μm) and an indium gallium phosphide (InGaP) photodetector (size: 185 μm × 125 μm × 3 μm), on a copper-clad PI film (Cu/PI/Cu: 18/25/18 μm). Contacts of the LED and the detector are metallized with sputtered metal layers (Cr/Au/Cu/Au = 6 nm/100 nm/600 nm/150 nm).

#### Fabrication of InGaP detectors

The InGaP detector structure is grown on gallium arsenide (GaAs) substrates via metal organic chemical vapor deposition (MOCVD). The epitaxially grown stack include (from top to bottom): a 500 nm n-type (Si-doped, 2×10^18^ cm^−3^) (Al_0.7_Ga_0.3_)_0.5_In_0.5_P filter layer, a 10 nm n-type GaAs (Si-doped, >6×10^18^ cm^−3^) top contact layer, a 30 nm n-type (Si-doped, 5×10^18^ cm^−3^) In_0.5_Al_0.25_Ga_0.25_P window layer, a 100 nm n-type (Si-doped, 2×10^18^ cm^−3^) In_0.5_Ga_0.5_P emitter layer, a 1 µm p-type (Zn-doped, 3×10^16^ cm^−3^) In_0.5_Ga_0.5_P base layer, a 100 nm p-type (Mg-doped, 2×10^18^ cm^−3^) In_0.5_Al_0.25_Ga_0.25_P back-surface field (BSF) layer, a 100 nm p-type (Mg-doped, >5×10^18^ cm^−3^) GaAs contact layer, a 1 µm p-type In_0.5_Ga_0.5_P (Mg-doped, 2×10^18^ cm^−3^) support layer and a 500 nm undoped Al_0.95_Ga_0.05_As sacrificial layer on a GaAs substrate (Table S1).

A detailed description of the process to fabricate the freestanding microscale photodetector is listed below:

1. Clean the epi-wafer with acetone, isopropyl alcohol (IPA), and deinoized (DI) water.
2. Dehydrate at 110 °C for 15 min.
3. Spin coat positive photoresist (PR) (SPR220-v3.0, Microchem, 500 rpm / 6 s, 3000 rpm / 30 s) and soft-bake at 110 °C for 1.5 min.
4. Expose PR with a UV lithography tool (URE-2000/25, IOE CAS) with a dose of 300 mJ/cm^2^ through a chrome mask, post-bake at 110 °C for 1.5 min.
5. Develop PR in an aqueous base developer (AZ300 MIF), rinse with DI water and hard-bake at 110 °C for 20 min.
6. Etch the (Al_0.7_Ga_0.3_)_0.5_In_0.5_P filter with a mixture of H_3_PO_4_ and HCl (1:1 by volume) for ∼7 s and rinse with DI water.
7. Etch the n-type GaAs contact and the In_0.5_Al_0.25_Ga_0.25_P window with a mixture of H_3_PO_4_, H_2_O_2_, and H_2_O (3:1:25 by volume) for ∼12 s and rinse with DI water.
8. Remove the PR using acetone, IPA and DI water.
9. Dehydrate at 110 °C for 15 min.
10. Spin coat positive photoresist (PR) (SPR220-v3.0, Microchem, 500 rpm / 6 s, 3000 rpm / 30 s) and soft-bake at 110 °C for 1.5 min.
11. Expose PR with UV lithography tools (URE-2000/25, IOE CAS) with a dose of 300 mJ/cm^2^ through a chrome mask and post-bake at 110 °C for 1.5 min.
12. Develop PR in aqueous base developer (AZ300 MIF), rinse with DI water and hard bake at 110 °C for 20 min.
13. Etch the In_0.5_Ga_0.5_P emitter, the In_0.5_Ga_0.5_P base and the In_0.5_Al_0.25_Ga_0.25_P BSF with concentrated HCl for ∼20 s.
14. Remove PR using acetone, IPA, DI water.
15. Dehydrate at 110 °C for 15 min.
16. Spin coat negative photoresist (AZ® nLOF 2070, MicroChemicals, 500 rpm / 6 s, 3000 rpm / 30 s) and soft-bake at 110 °C for 2 min.
17. Expose PR with UV lithography tools (URE-2000/25, IOE CAS) with a dose of 60 mJ/cm^2^ through a chrome mask and post-bake at 110 °C for 75 s.
18. Develop PR in aqueous base developer (AZ300 MIF), rinse with DI water.
19. Sputter 6 nm/200 nm of Cr/Au.
20. Lift-off in acetone, and clean the wafer using acetone, IPA, DI water.
21. Dehydrate at 110 °C for 15 min.
22. Pattern PR (SPR220-3.0, Microchem).
23. Etch the p+ GaAs contact in a mixture of H_3_PO_4_/H_2_O_2_/H_2_O (3:1:25, by volume) for 40 s and rinse with DI water.
24. Etch the In_0.5_Ga_0.5_P support and the Al_0.95_Ga_0.05_As sacrificial layer with concentrated HCl for ∼40 s.
25. Remove the PR using acetone, IPA, DI water.
26. Pattern PR (SPR220-3.0, Microchem).
27. Etch in diluted HF (49% HF : DI water = 1:10, by volume) to release the devices from GaAs growth substrates for ∼2.5 h and rinse with DI water.
28. The free-standing InGaP photodetector arrays are formed and ready for picking up.

#### Fabrication of violet InGaN LEDs

The InGaN based violet LED structures are grown on 2-inch planar sapphire substrates using MOCVD. The main LED structure (from bottom to top) includes: the sapphire substrate, a GaN buffer layer, an n-GaN, an InGaN / GaN multiple-quantum-well (MQW) layer, a p-GaN and an indium tin oxide (ITO) layer for top contacts. The thickness of the entire epitaxial structure is about 10 μm. LED devices are lithographically fabricated, with Cr/Au layers serving as ohmic contacts. Inductively couple plasma reactive ion etching (ICP-RIE) is used to define the LED mesa (lateral dimension 190 μm × 110 μm). After bonding the fully fabricated LED arrays onto a thermal release tape (TRT) (Nitto Denko Corp.), laser lift-off (LLO) is applied to separate the thin-film LEDs from sapphire substrates by thermally decompose GaN into gallium (Ga) metal and nitrogen gas at the interface. A krypton fluoride (KrF) excimer laser at 248 nm (Coherent, Inc., CompexPro110) serves as the light source, with a uniform irradiation area of 5 mm × 15 mm. The power density during LLO is optimized to be around 0.6 J/cm^2^ for LEDs. After laser irradiation, the micro-LEDs are released from sapphire by mild mechanical force at 70 °C (melting point of Ga is 29.7 °C). By heating up to 120 °C, the LEDs are detached from the TRT (the critical release temperature is about 110 °C) and ready for transfer printing by PDMS.

#### Fabrication of thin-film filters

The filter structure is deposited on a GaAs substrate using ion beam-assisted sputter deposition. The multilayered structure consists of titanium dioxide (TiO_2_) and silicon dioxide (SiO_2_) films with a total thickness of about 6.6 μm. Its detailed layer structure can be found in our previous work^1^. Its shape (size: 220 μm × 150 μm) is defined through femtosecond laser-cutting (800 nm, Ti-sapphire laser system, Coherent Inc.). Subsequently, the GaAs substrate is removed in a mixed solution with NH_4_OH, H_2_O_2_ and H_2_O (1:1:2, by volume) for ∼8 hours. Finally, PDMS stamps (Sylgard 184 from Dow Corning) are utilized to transfer and print the freestanding thin-film filters onto other foreign substrates.

#### Synthesis of the PtTFPP/PDMS film

To fabricate the oxygen-sensitive film, commercial platinum(II)-5,10,15,20-tetrakis (2,3,4,5,6-pentafluorophenyl)-porphyrin (PtTFPP) powders (PtT975, Frontier Scientific) are dissolved in toluene (0.08 mol/L), followed by the addition of the base component (part A) of polydimethylsiloxane (PDMS) to the solution, with different PtTFPP: PDMS weight ratios (0–0.50 wt%). After evaporating the solvent overnight, the substance is mixed with curing agent (part B) of PDMS with a weight ratio of 10:1 (part A/part B). Subsequently, vacuum evacuation removes air bubbles. The PtTFPP/PDMS film is ready for dip coating on the optoelectronic micro-probe.

#### Preparation of the adhesive solution

The adhesive solution comprises: bisphenol A glycerolate (1 glycerol/phenol) diacrylate, 3-(Trimethoxysilyl) propyl methacrylate, spin-on-glass (SOG 500F, Filmtronics Inc.), 2-Benzyl-2-(dimethylamino)-4’-morpholinobutyrophenone, and anhydrous ethanol^2^. The weight ratio is 200:100:100:9:2000. Stir at room temperature until full mixing. Store the mixed solution in refrigerator (4 °C) for future use.

### Fabrication of the microscale optoelectronic probe

A detailed description of the fabrication process is listed below:

#### Substrate preparation

1. Cut the copper-clad PI substrate (Cu/PI/Cu: 18/25/18 μm, DuPont) into small pieces (size: 15 mm × 18 mm).
2. Laminate the copper-clad polyimide film onto a PDMS coated glass slide

a. Degas PDMS (Sylgard 184) 10 : 1 (base: curing agent, by weight).
b. Spin cast PDMS to a glass slide at 500 rpm/6 s, 3000 rpm/30 s to form a PDMS film (thickness: ∼10 μm).
c. Bake the glass for 23 s at 110 °C.
d. Laminate the PI film onto the PDMS coated glass. Make sure that the polyimide attaches to PDMS tightly without any bubbles.
e. Bake the sample for 30 mins at 110 °C until PDMS is fully cured.
f. Clean the sample by acetone, IPA and DI water.
g. Dehydrate at 110 °C for 15 min.
h. Spin cast 8 μm polyimide (YDPI-102, Jing’ai Microelectronics) at 500 rpm/9 s, 3000 rpm/30 s to form an insulating film.
i. Bake the sample for 1 h at 80 °C, 1 h at 130 °C, 1 h at 180 °C, 3 h at 230 °C.

#### InGaP detector transfer

3. Clean the PI sample by acetone, IPA and DI water.
4. Dehydrate at 110 °C for 15 min.
5. Spin coat negative photoresist (PR) (AZ nLOF 2070, 500 rpm / 6 s, 3000 rpm / 30 s) and soft-bake at 110 °C for 2 minutes 20 seconds.
6. Expose PR with UV lithography tools (URE-2000/25, IOE CAS) with a dose of 60 mJ/cm2 (URE-2000/25, IOE CAS) through a chrome mask and post exposure bake at 110 °C for 75 s.
7. Develop PR in an aqueous base developer (AZ300 MIF) and rinse with DI water.
8. Deposit 50 nm Al as the markers by sputtering.
9. Lift off PR in acetone.
10. Clean the substrate and dehydrate at 110 °C for 10 min.
11. Spin coat the prepared adhesive solution (500 rpm/6 s, 3000 rpm/30 s) on the substrate and soft-bake at 110 °C for 3 mins.
12. Transfer print InGaP detectors from the source GaAs wafer onto the processed substrate with PDMS stamps.
13. Cure under UV for 10 minutes and bake at 110 °C for 3 h.
14. Clean1 the PR by reactive ion etching (RIE) (O2, 200 sccm, 90 mTorr, 150 W) for 12 min.

#### InGaP detector encapsulation

15. Clean the sample (acetone, IPA, DI water) and dehydrate at 110 °C for 15 min.
16. Clean by RIE (O2, 200 sccm, 90 mTorr, 150 W) for 1 min.
17. Spin coat SU8-3005 epoxy (500 rpm/6 s, 3000 rpm/30 s).
18. Soft-bake at 65 °C for 1 min and 95 °C for 3 min.
19. Pattern SU8 to expose contact pads of the detector by UV lithography with a dose of 210 mJ/cm2.
20. Post-bake at 65 °C for 1 min and 95 °C for 2 min.
21. Develop in propylene glycol monomethyl ether acetate (PGMEA) for 2.5 min and rinse with IPA.
22. Hard bake at 130 °C for 2h.

#### InGaP detector metallization

23. Clean the sample (acetone, IPA, DI water) and dehydrate at 110 °C for 15 min.
24. Pattern PR AZ nLOF 2070.
25. Deposit Cr/Au/Cu/Au = 6 nm/100 nm/600 nm/150 nm by sputtering.
26. Lift-off PR in acetone.

#### Filter transfer

27. Clean the sample (acetone, IPA, DI water) and dehydrate at 110 °C for 15 min.
28. Spin coat 2 μm polyimide (500 rpm/6 s, 3000 rpm/30 s) and bake at 110 °C for 2 h.
29. Spin coat 2 μm polyimide (500 rpm/6 s, 3000 rpm/30 s).
30. Bake at 85 °C for 10 s.
31. Transfer print dielectric filters on top of detectors with PDMS stamps.
32. Hard-bake at 110 °C for 2 h.

#### InGaN LED transfer

1. 33. Clean the sample (acetone, IPA, DI water) and dehydrate at 110 °C for 15 min.
2. 34. Spin coat 8 μm polyimide (500 rpm/6 s, 3000 rpm/30 s).
3. 35. Bake at 85 °C for 2 min.
4. 36. Transfer print InGaN violet LEDs onto the substrate with PDMS stamps.
5. 37. Hard-bake at 85 °C for 1 h and 130 °C for 4 h.

#### InGaN LED encapsulation

38. Clean the sample (acetone, IPA, DI water) and dehydrate at 110 °C for 15 min.
39. Clean by RIE (O2, 200 sccm, 90 mTorr, 150 W) for 1 min.
40. Spin coat SU8-3005 epoxy (500 rpm/6 s, 3000 rpm/30 s).
41. Soft-bake at 65 °C for 1 min and 95 °C for 3 min.
42. Pattern SU8 to expose contact pads of the LED by UV lithography with a dose of 210 mJ/cm2.
43. Post-bake at 65 °C for 1 min and 95 °C for 2 min.
44. Develop in PGMEA for 2.5 min and rinse with IPA.
45. Hard bake at 130 °C for 2h.

#### InGaN LED metallization

46. Clean the sample (acetone, IPA, DI water) and dehydrate at 110 °C for 15 min.
47. Pattern PR AZ nLOF 2070.
48. Deposit Cr/Au/Cu/Au = 6 nm/100 nm/600 nm/150 nm by sputtering.
49. Lift-off PR in acetone.

#### Device encapsulation

50. Clean the sample (acetone, IPA, DI water) and dehydrate at 110 °C for 15 min.
51. Expose under ultraviolet induced ozone (UV Ozone, PCE-22-LD, YIMA) for 15 min.
52. Pattern SU8-3005 epoxy (500 rpm / 5 s, 3000 rpm / 30 s), cure under UV for 210 mJ / cm2 and bake at 110 °C for 30 min.

#### Laser milling

53. Use laser milling to cut the PI film into micro-probe shapes (LPKF ProtoLaser U4, 365 nm; 1000 Hz, 0.5 W, 50 repeats + 1 W, 100 repeats + 2W, 40 repeats)

#### Device encapsulation

54. Expose the micro-probes under UV ozone for 15 min.
55. Encapsulate the micro-probes with PDMS

a. Degas PDMS 10:1 (base: curing agent, by weight).
b. Dip coat the micro-probes with PDMS.
c. Bake the probes in 75 °C for 2 h.
56. Coat the probes with a layer of parylene C (14 μm, Specialty Coating Systems).
57. Clean the probes with RIE (O2, 200 sccm, 90 mTorr, 150 W) for 1 min.
58. Encapsulate the probes with another layer of PDMS.
59. Dip the synthesized PtTFPP/PDMS on the tip of the probe.

---

**Figure S1.**
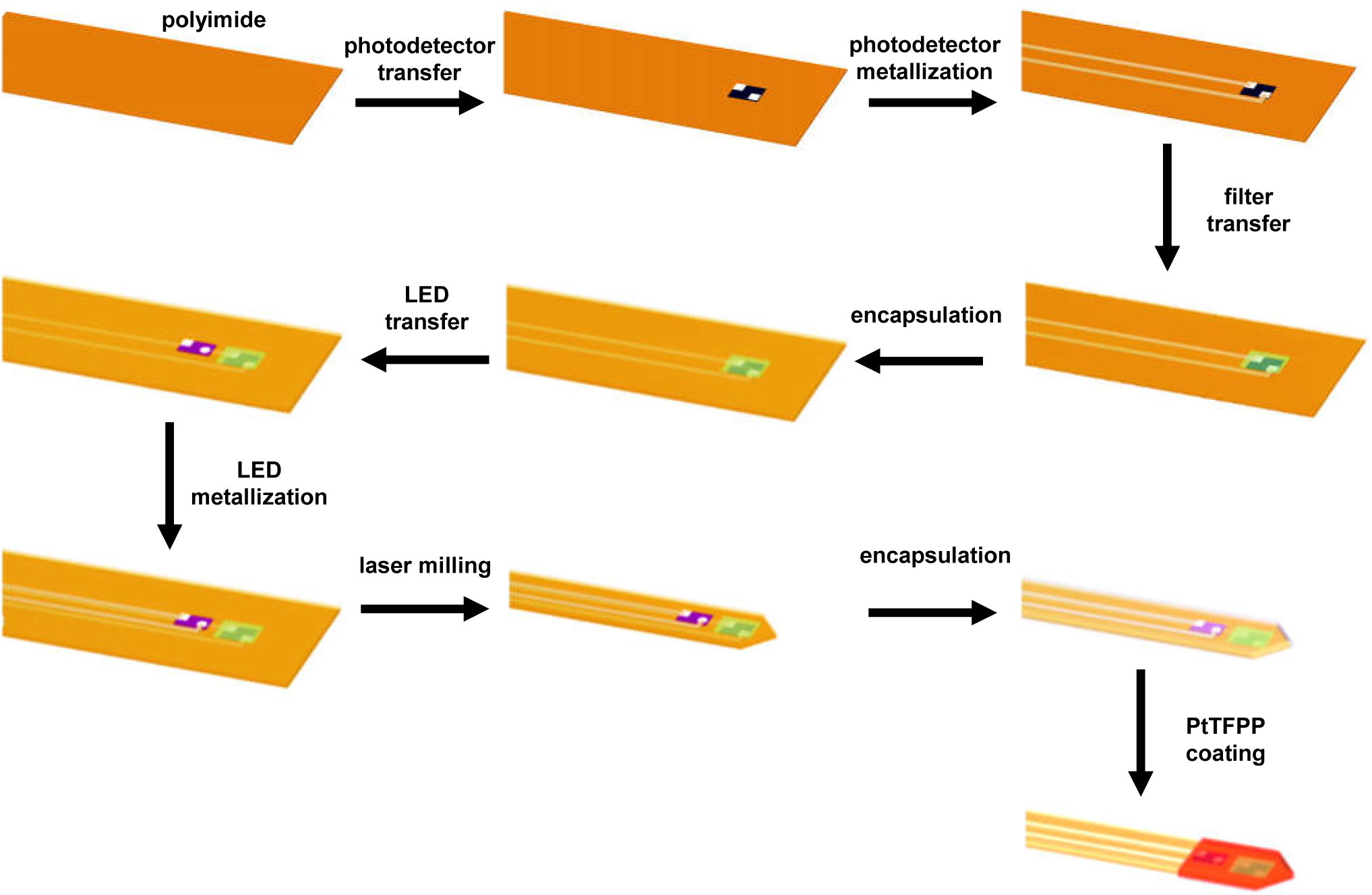
Schematic illustration of the process flow for fabricating the PbtO_2_ sensing probe.

**Figure S2.**
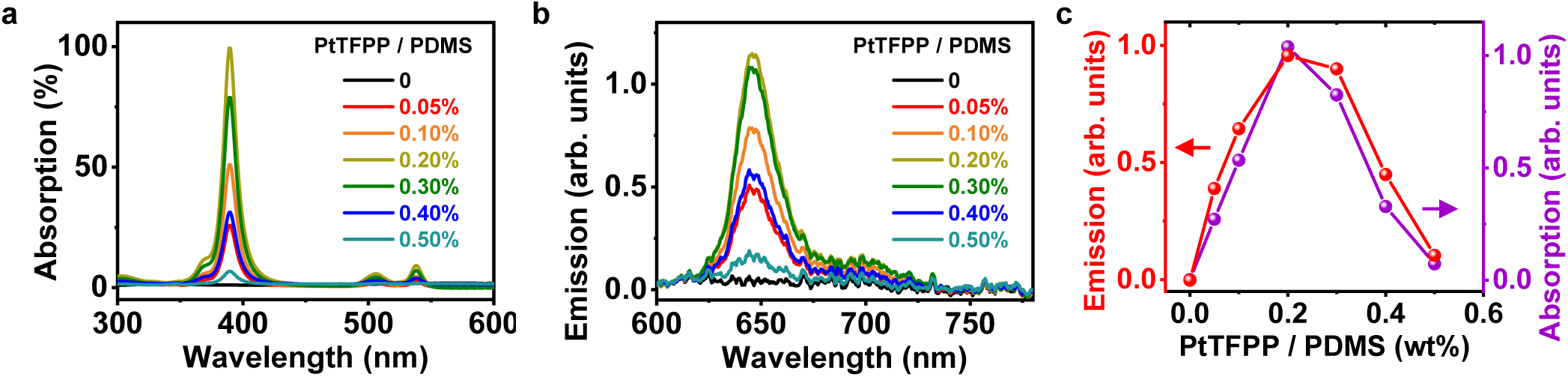
Optical properties of PtTFPP/PDMS films. (a) Absorption and (b) PL emission spectra of PtTFPP with different mass concentrations mixed in PDMS films (thickness 10 μm). (c) Peak emissions and absorptions as a function of PtTFPP mass concentrations.

**Figure S3.**
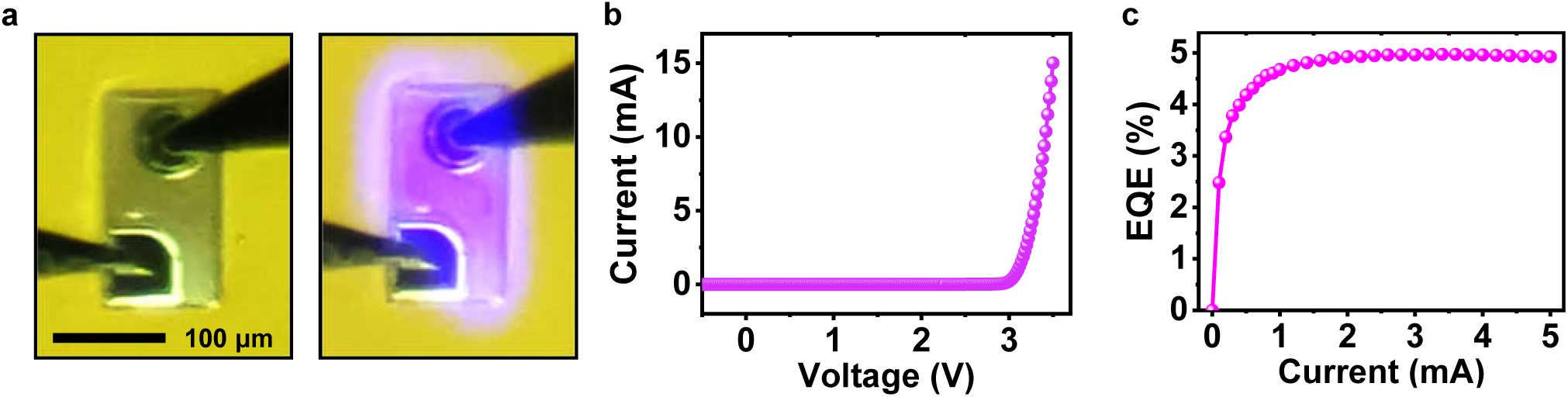
Images and optoelectronic properties of the InGaN violet LED. (a) Microscopic images of a violet LED with (right) and without (left) electroluminescence. (b) The LED’s current– voltage characteristic curve. (c) The LED’s external quantum efficiencies (EQEs) as a function of currents.

**Figure S4.**
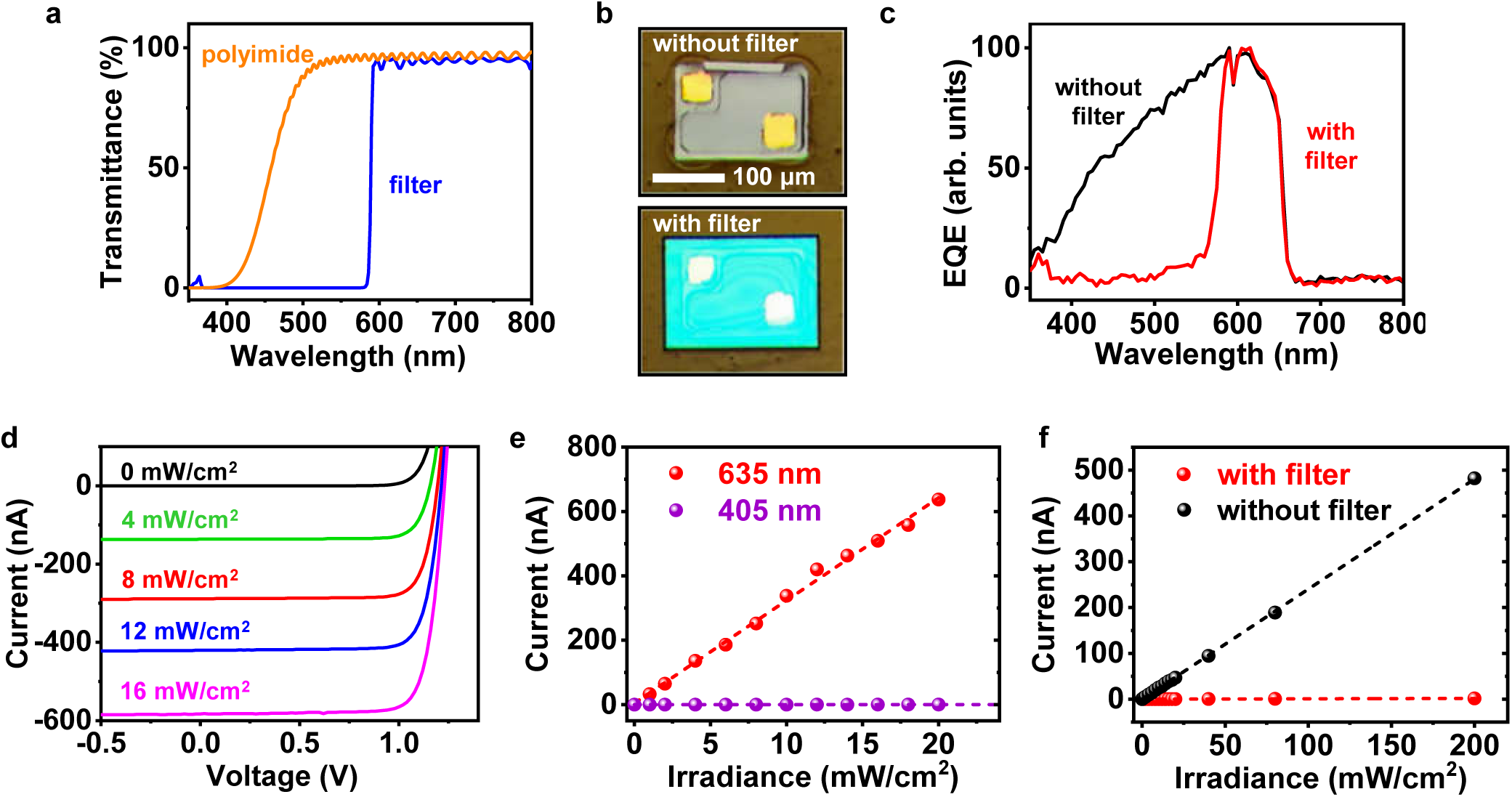
Images and optoelectronic properties of the InGaP photodetector. (a) Transmission spectra of a polyimide film (thickness 8 μm) and a dielectric filter. (b) Images of InGaP detectors with (lower) and without (upper) the filter. (c) External quantum efficiency (EQE) spectra for InGaP detectors with and without coatings of polyimide and the filter. (d) Current–voltage curves of an InGaP detector under different irradiance at 635 nm. (e) Photocurrent response of a detector coated with a filter measured at 405 nm and 635 nm illuminations. (f) Photocurrent response of a detector with and without filter coatings under 405 nm illumination.

**Figure S5.**
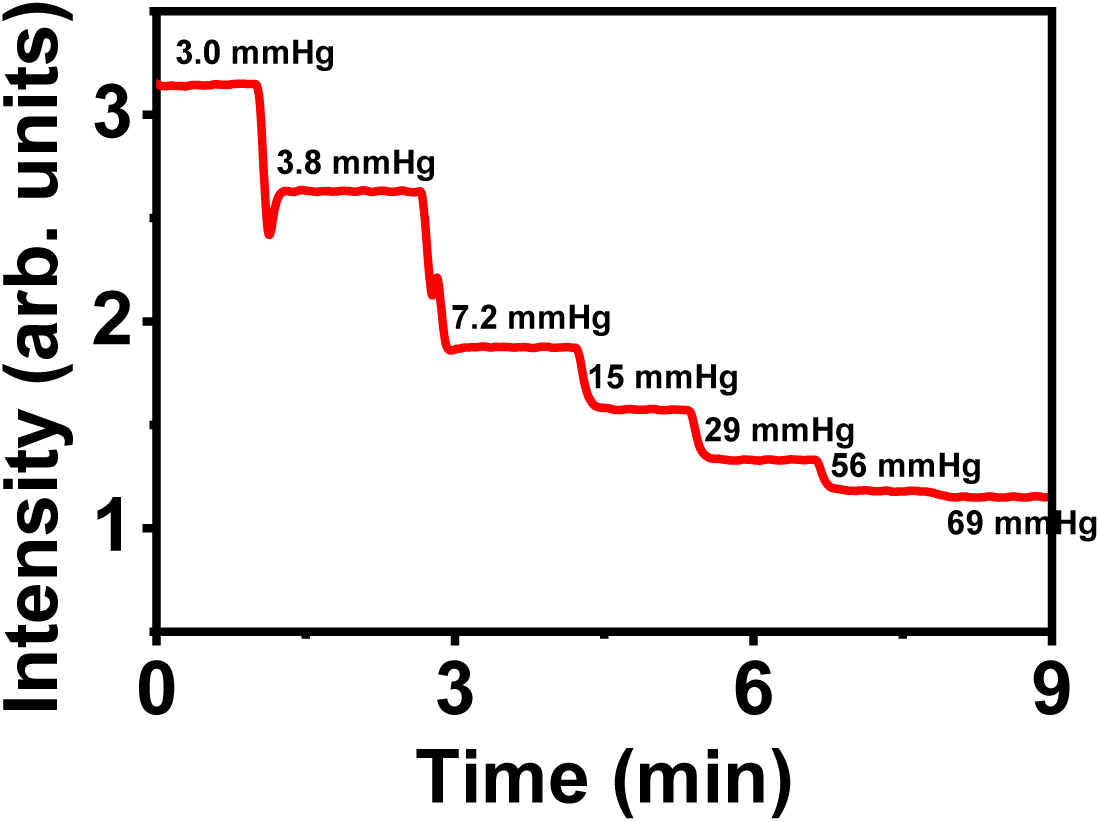
Detection limit of the probe. Continuously monitored PL intensities with the probe at different O_2_ partial pressures (pO_2_). The chamber pressure is kept at 1 atm (∼760 mmHg). The ratios of N_2_ and O_2_ gas flows (unit: L/min) are varied: 25/0.1, 20/0.1, 10/0.1, 10/0.2, 10/0.4, 10/0.8 and 10/1. Corresponding pO_2_ are 3.0, 3.8, 7.2, 15, 29, 56, and 69 mmHg.

**Figure S6.**
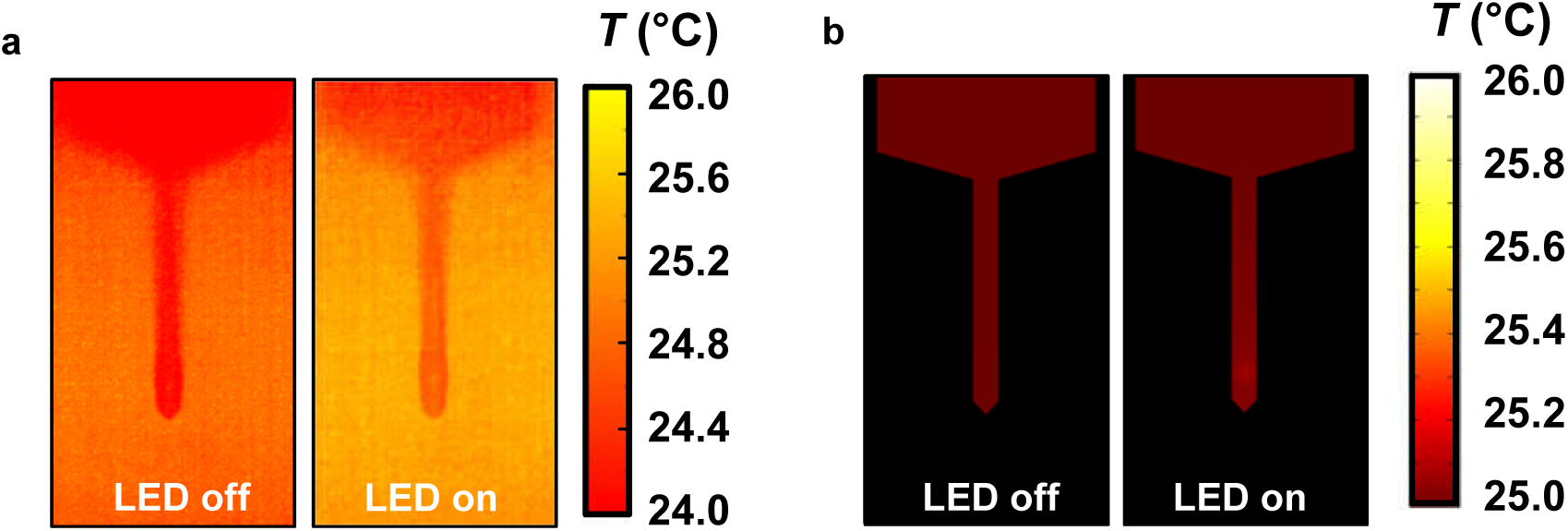
Thermal properties of the probe. (a) Measured and (b) simulated temperature distributions on the probe surface when the violet LED is off (left) or on (right) in air (LED current 0.5 mA, frequency 4 Hz, duty cycle 5%).

**Figure S7.**
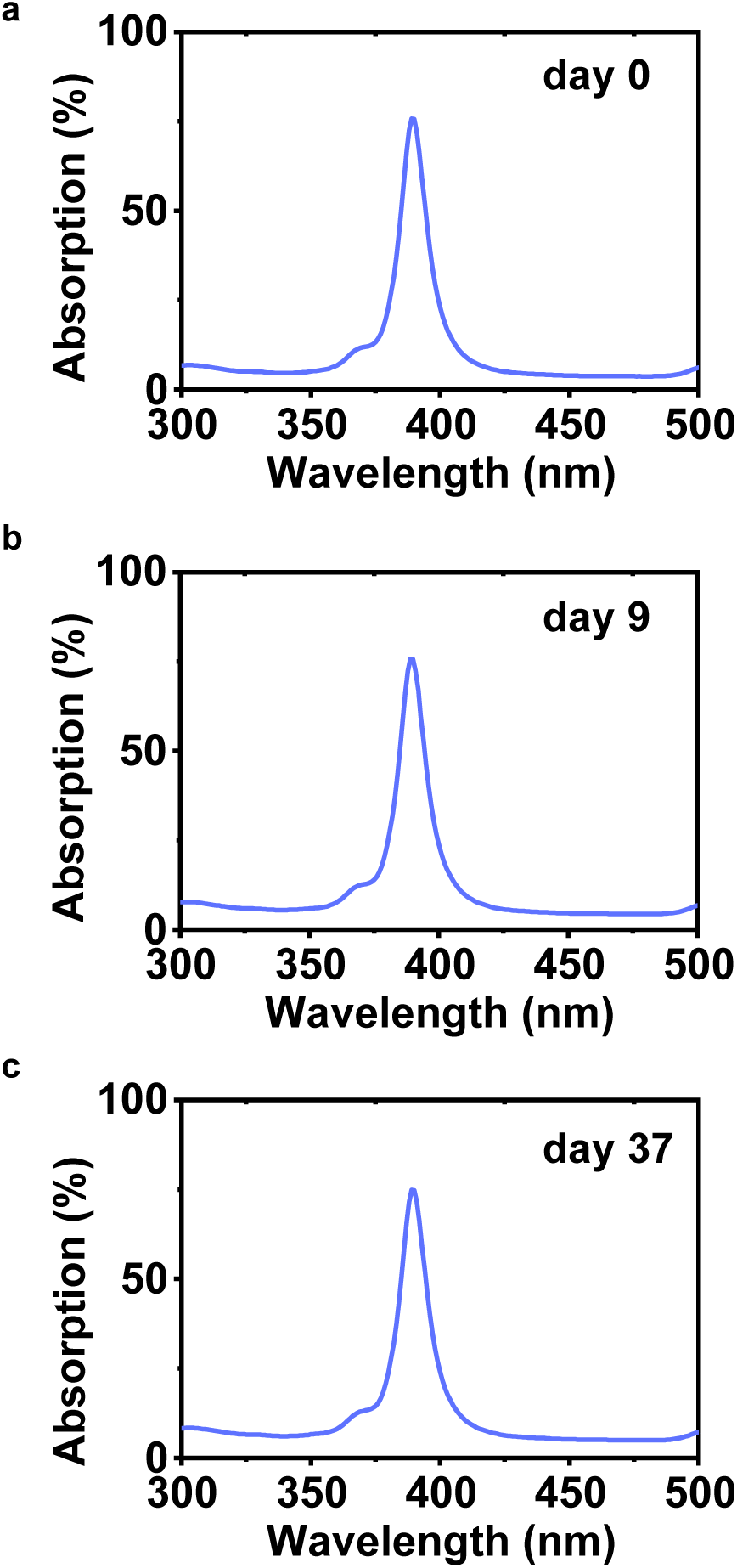
Absorption spectra of the oxygen sensing film immersing the film in PBS for 0 day (a), 9 days (b) and 37 days (c).

**Figure S8.**
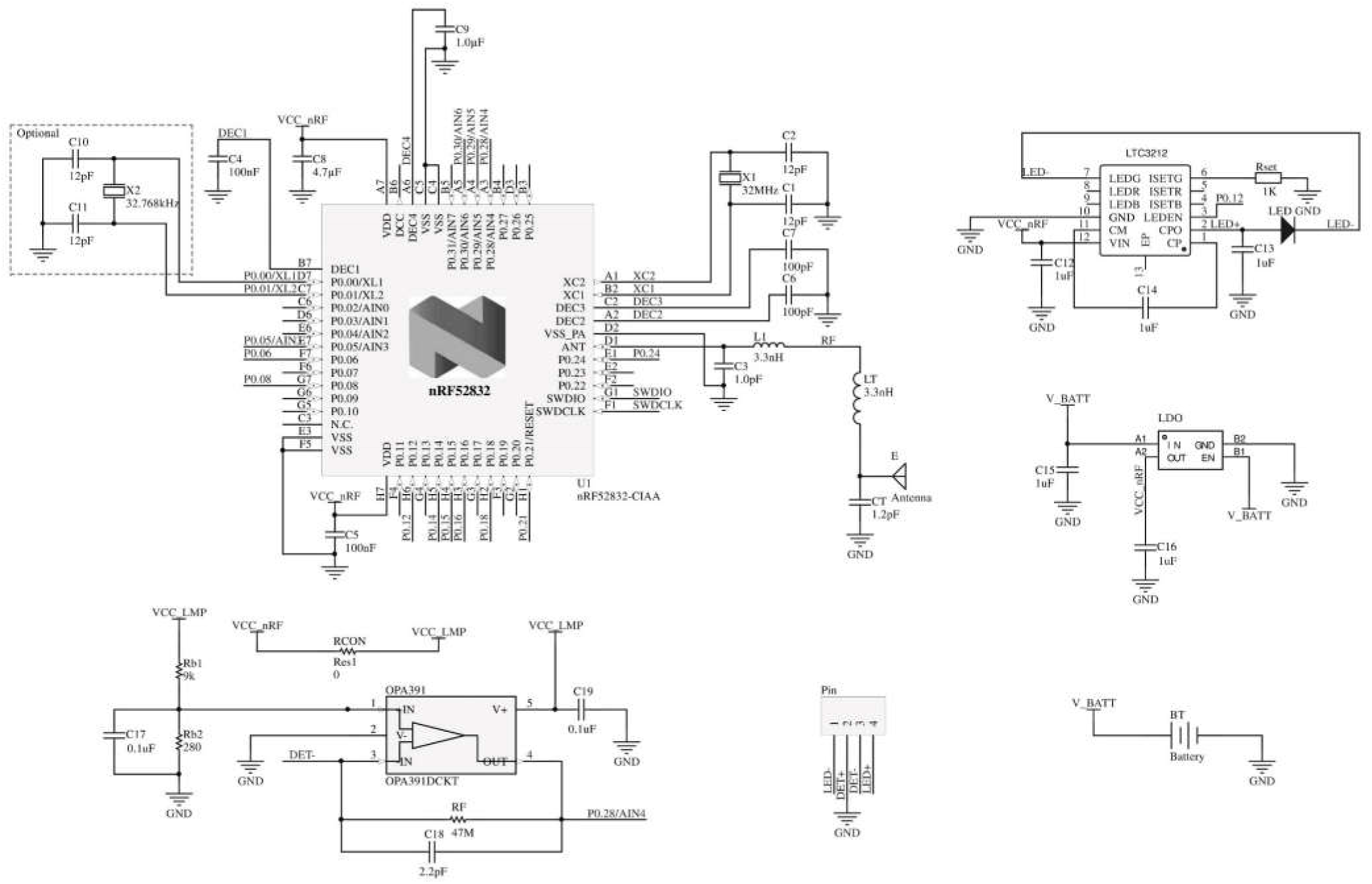
Detailed circuit design diagram of the wireless, battery-powered circuit system.

**Figure S9.**
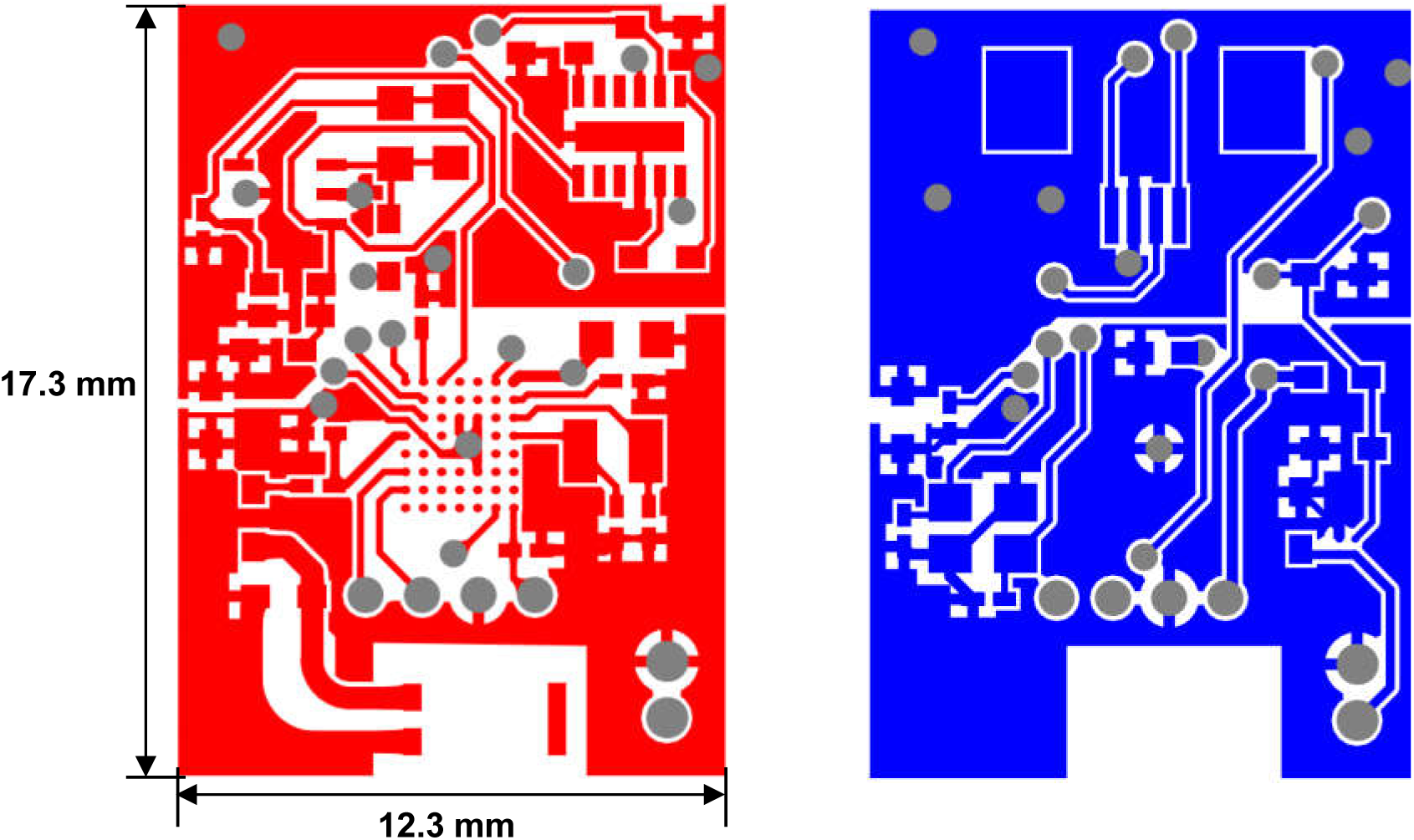
Layout of the designed printed circuit board for the wireless, battery-powered circuit system (red, top layer; blue, bottom layer).

**Figure S10.**
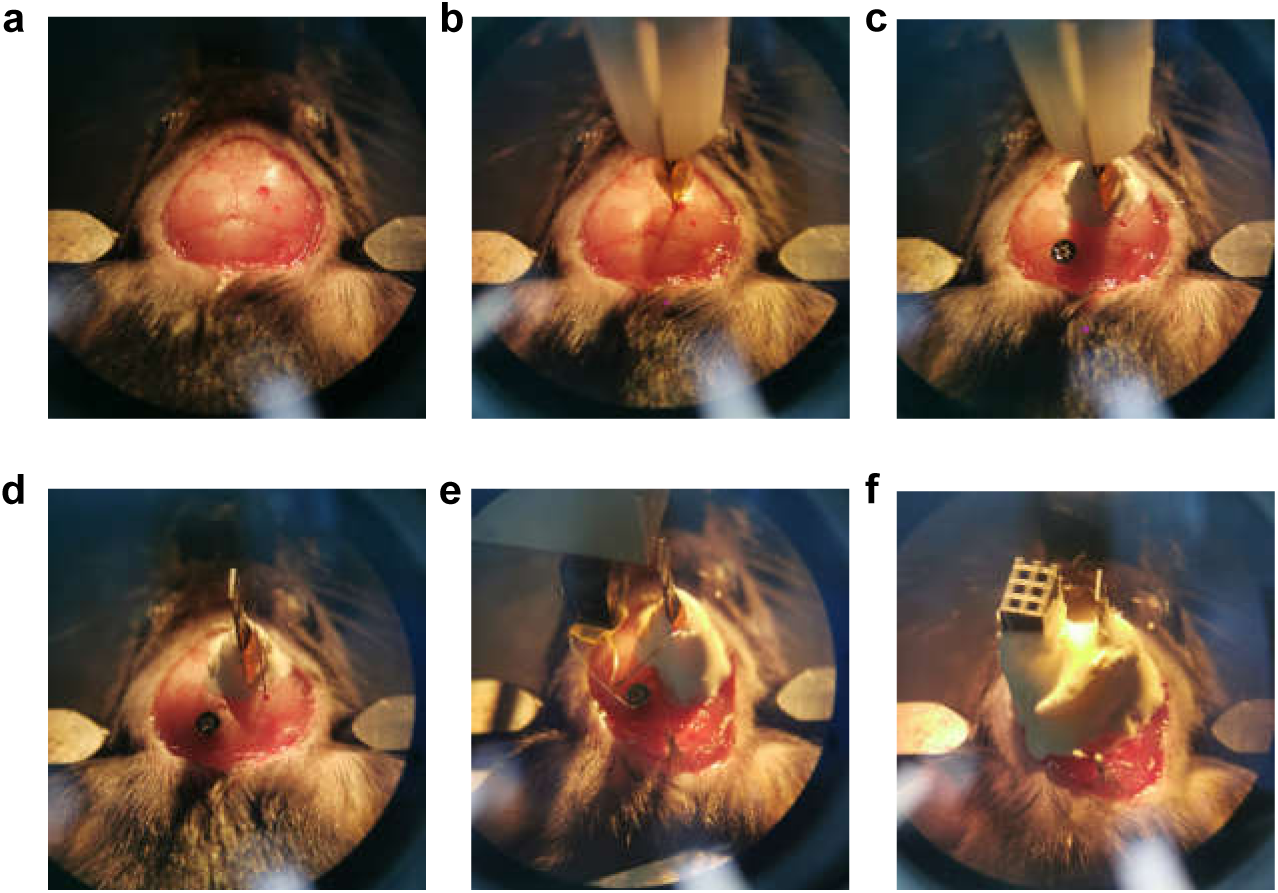
Photographs showing the surgical procedure of implanting a PbtO_2_ sensing probe as well as recording and stimulation electrodes into the mouse hippocampus. (a) Expose the skull, and drill a hole on the skull. (b) Implant the probe into mouse brain. (c) Apply dental cements to fix the probe to the skull, and implant a screw as a reference electrode. (d) Implant stimulation and recording electrodes into the brain. (e) Fix the electrodes with dental cements. (f) Wire the electrodes to a connector, and apply more dental cements to fix all components.

**Figure S11.**
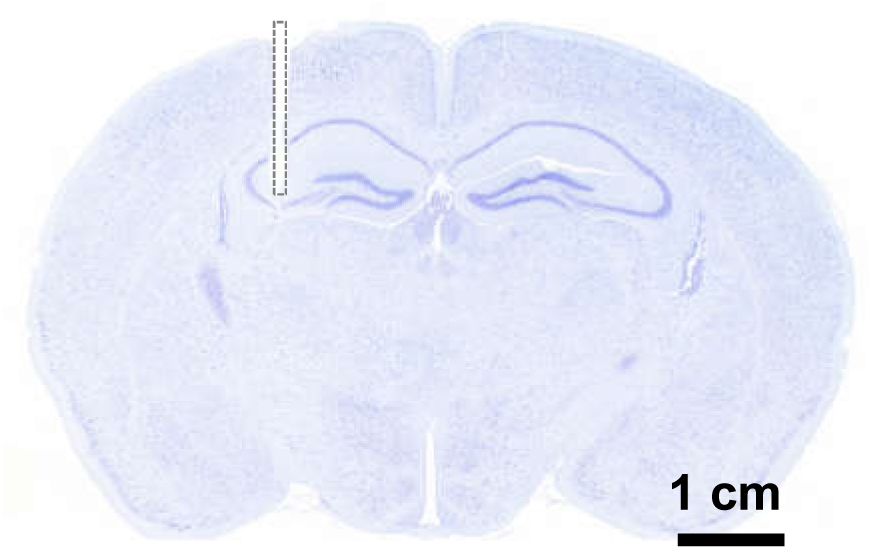
Coronal section stained with Nissl, showing the lesion region in the hippocampus (CA3) of mice created by the probe.

**Figure S12.**
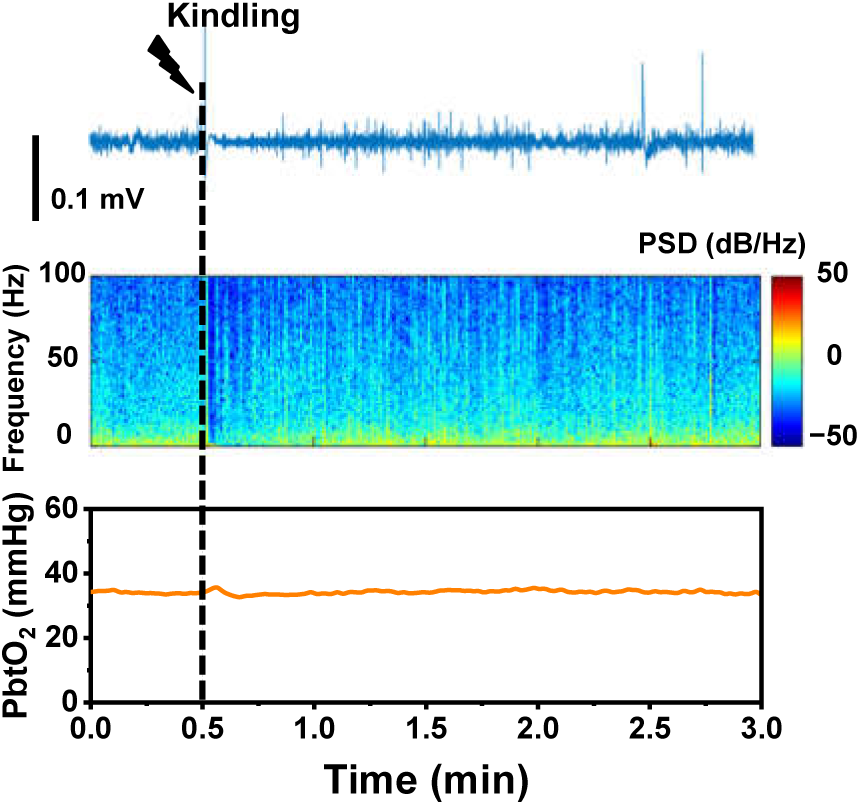
Simultaneously recorded electrophysiological activities in CA1 and PbtO_2_ in CA3 under kindling from a mouse without afterdischarge. Top: LFP trace. Middle: power spectral density (PSD) of LFP. Bottom: PbtO_2_.

**Figure S13.**
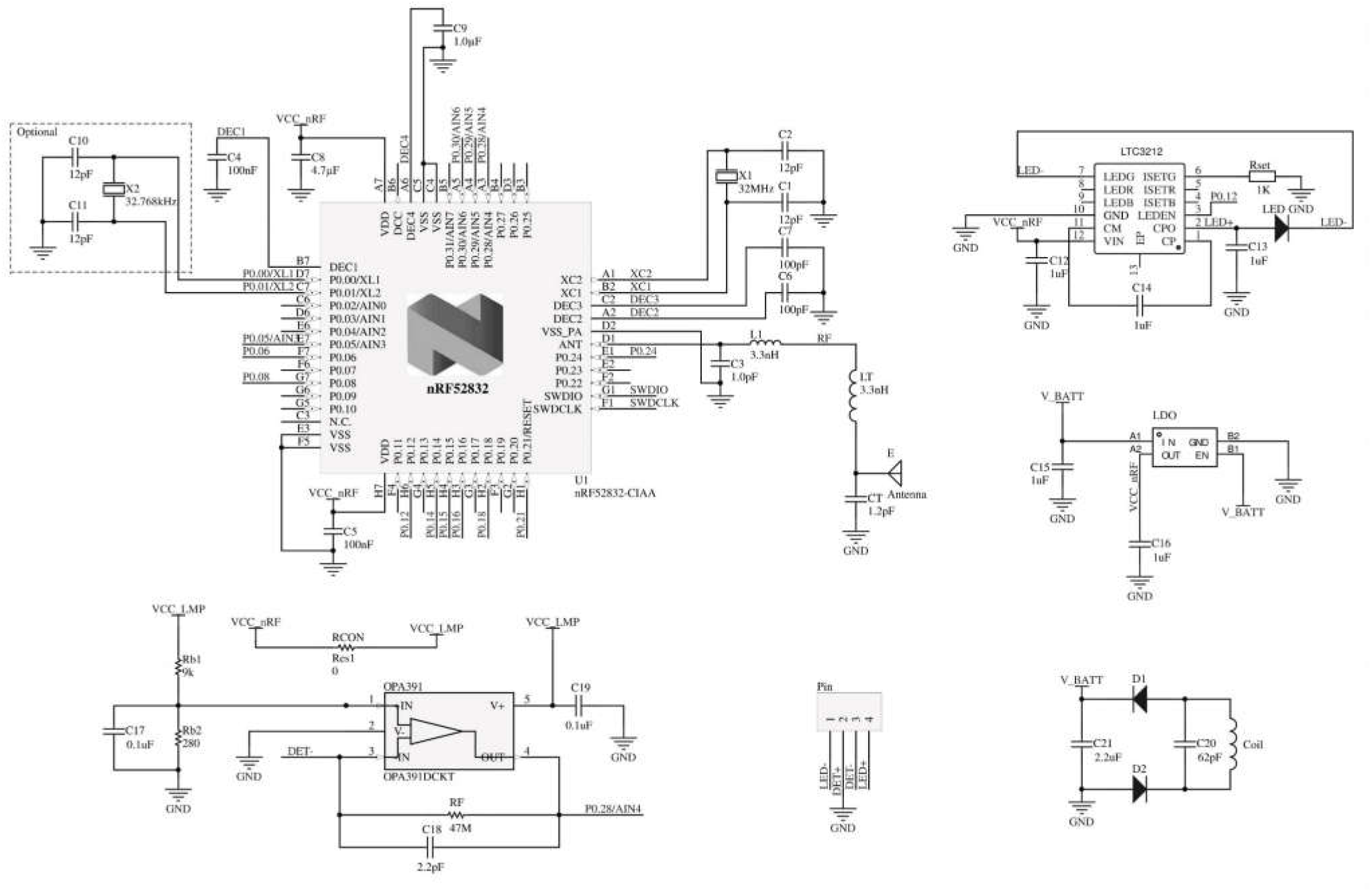
Detailed circuit design diagram of the wireless, battery-free circuit system.

**Figure S14.**
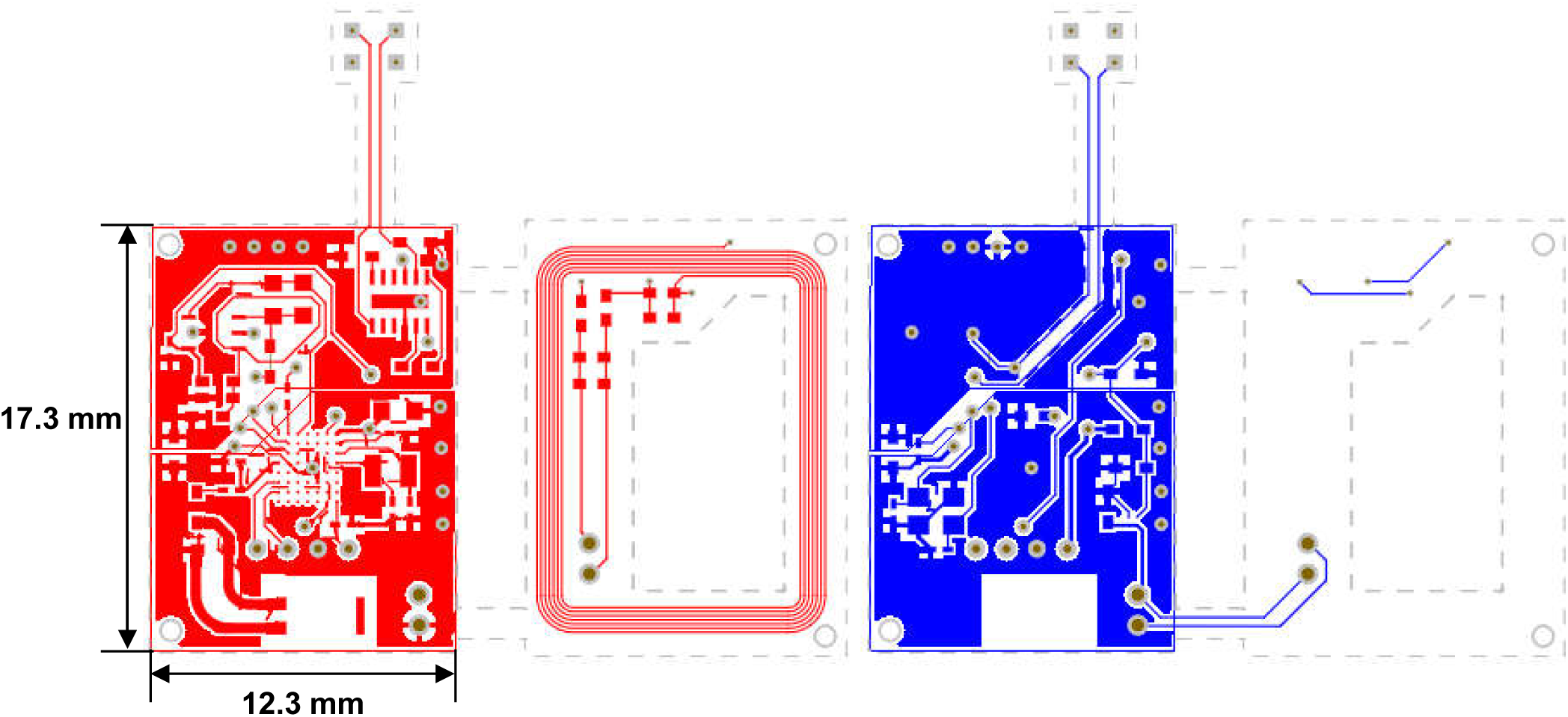
Layout of the designed printed battery-free circuit board for the wireless, battery-free circuit system (red, top layer; blue, bottom layer).

**Figure S15.**
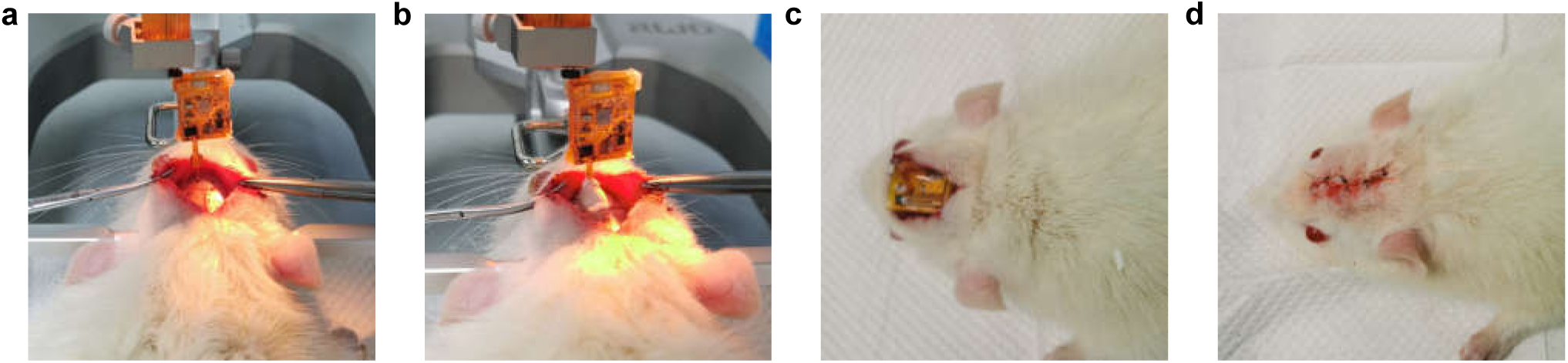
Photographs showing the surgical procedure of implanting a probe and its associated battery-free circuit in a rat. (a) Expose the skull, drill a hole on the skull and implant the probe into the rat brain. (b) Apply the dental cement to fix the probe to the skull. (c) Place the circuit underneath the skin. (d) Suture the skin.

**Figure S16.**
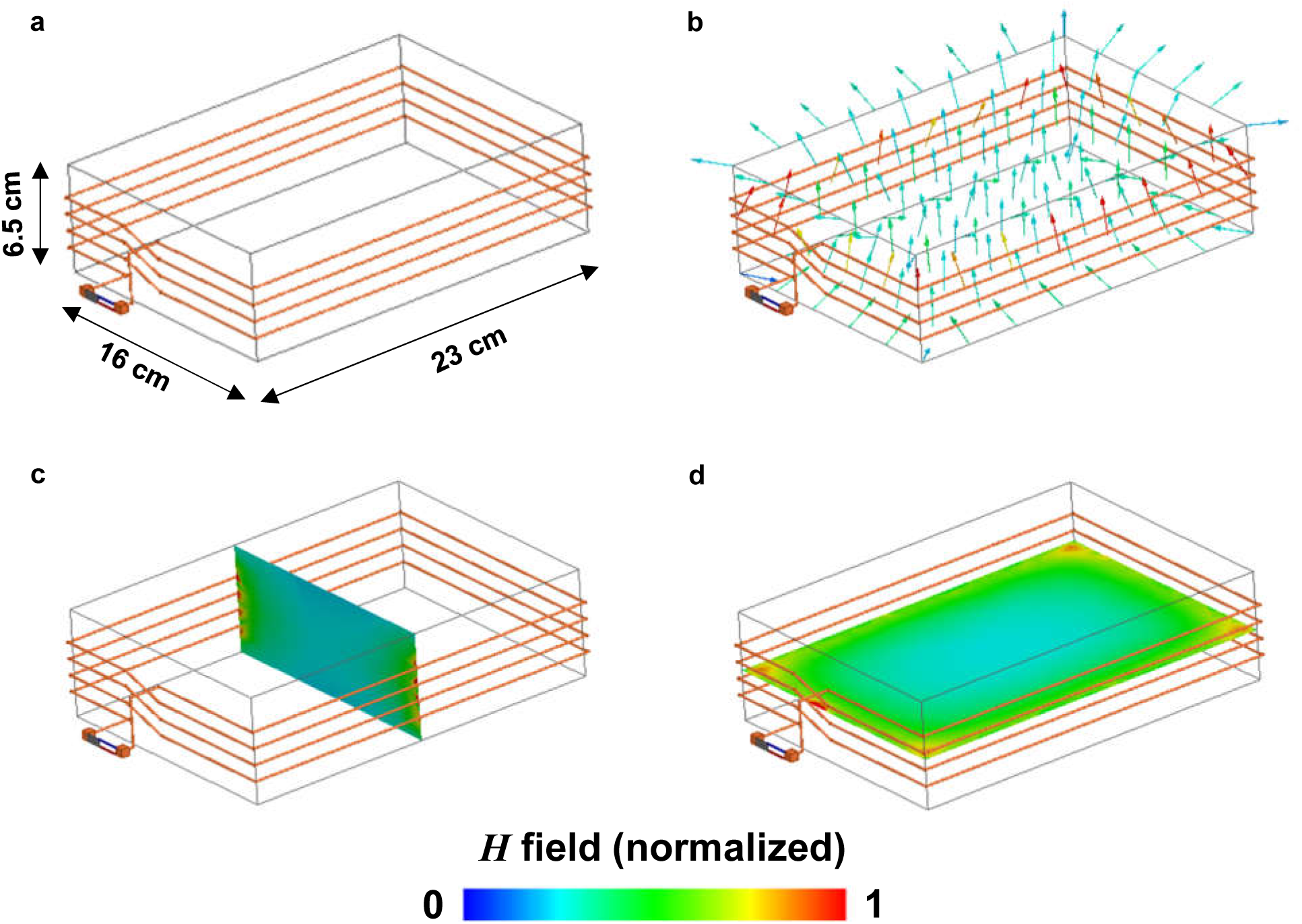
(a) Schematic illustration of the antenna system for wireless power transfer (dimension: *W* × *L* × *H* = 16 cm × 23 cm × 6.5 cm). (b) Simulated magnetic field distribution (normalized) inside the box. (c) Longitudinal and (d) Cross-sectional view of the field distribution.

**Figure S17.**
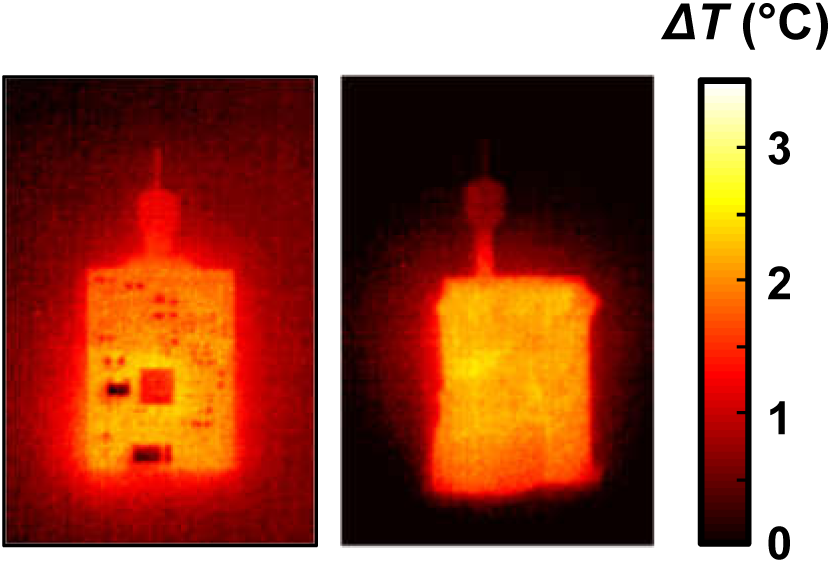
Thermal properties of wireless circuits. (a) Battery-powered circuit. (b) Inductively powered, battery-free circuit (coated with PDMS).

**Table S1.** Epitaxial structure of the InGaP based detector wafer.

| Materials | Thickness (nm) | Doping (cm<sup>-3</sup>) | Dopant |
| --- | --- | --- | --- |
| n+ In<sub>0.5</sub>Al<sub>0.7</sub>Ga<sub>0.3</sub>P filter | 500 | 2e18 | Si |
| n+ GaAs contact | 10 | >6e18 | Si |
| n+ In<sub>0.5</sub>Al<sub>0.25</sub>Ga<sub>0.25</sub>P window | 30 | 5e18 | Si |
| n+ In<sub>0.5</sub>Ga<sub>0.5</sub>P emitter | 100 | 2e18 | Si |
| p In<sub>0.5</sub>Ga<sub>0.5</sub>P base | 1000 | 3e16 | Zn |
| p+ In<sub>0.5</sub>Al<sub>0.25</sub>Ga<sub>0.25</sub>P BSF | 100 | 2e18 | Mg |
| p+ GaAs contact | 100 | >5e18 | Mg |
| p+ In<sub>0.5</sub>Ga<sub>0.5</sub>P support | 1000 | 2e18 | Mg |
| Al<sub>0.95</sub>Ga<sub>0.05</sub>As sacrificial | 500 | undoped | - |
| GaAs substrate | - | - | - |

**Table S2.**
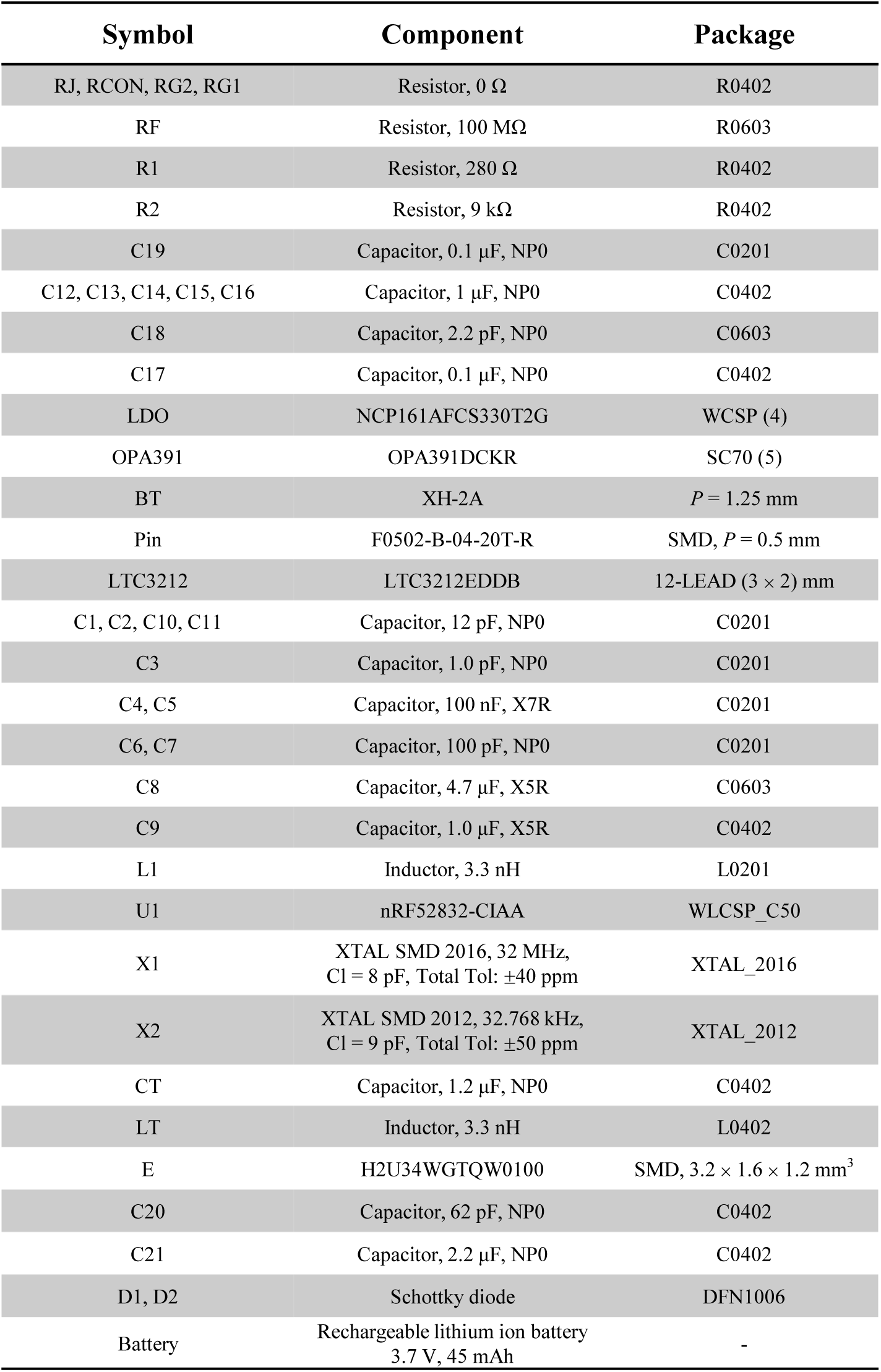
List of components used in the wireless circuit systems powered by a battery or an RF coil.

**Table S3.**
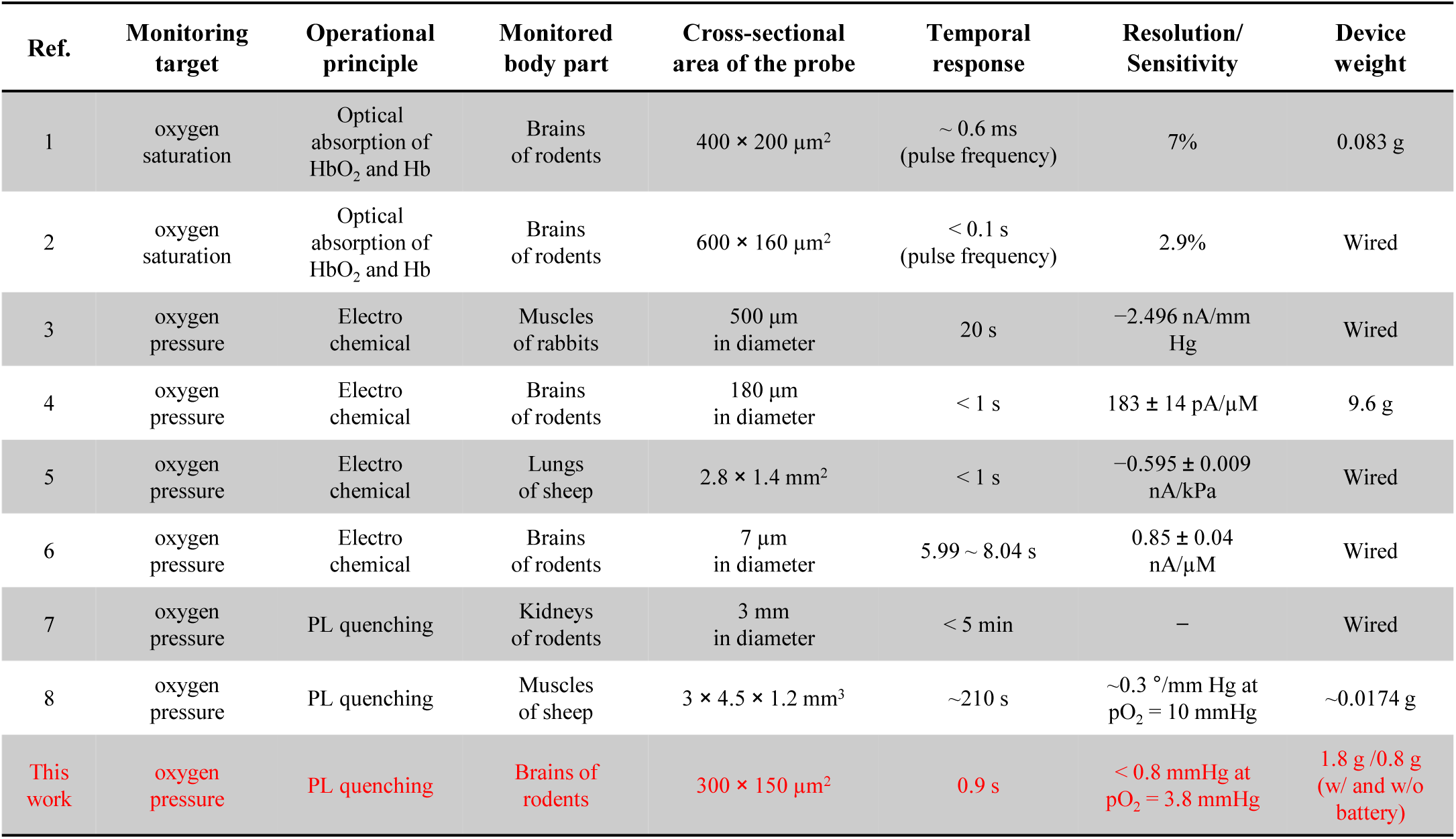
Comparing characteristic parameters of wireless and wired implantable oxygenation sensors reported in the literature and this work.

**Movie S1.**
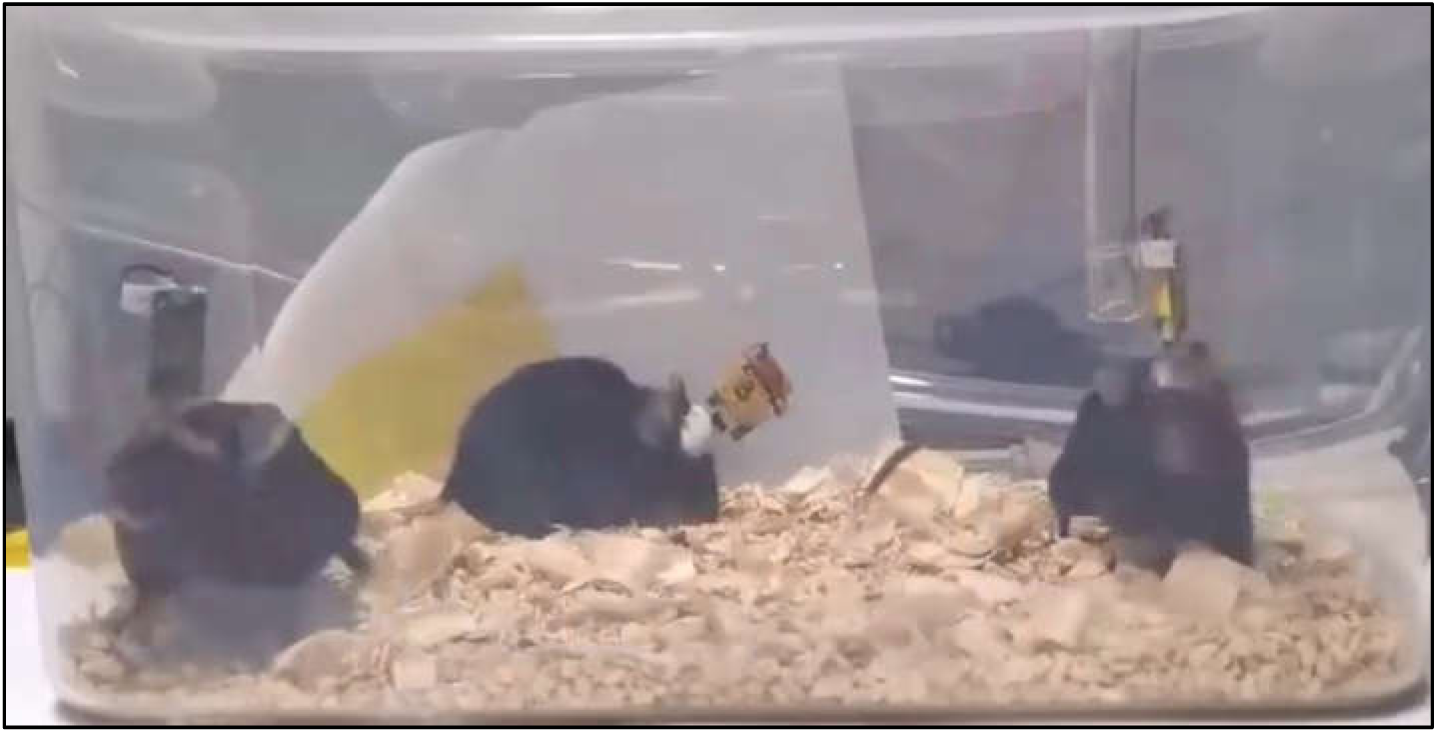
Video for three behaving mice implanted with oxygen sensing probes and head-mounted circuits. Their PbtO_2_ signals are recorded simultaneously.

**Movie S2.**
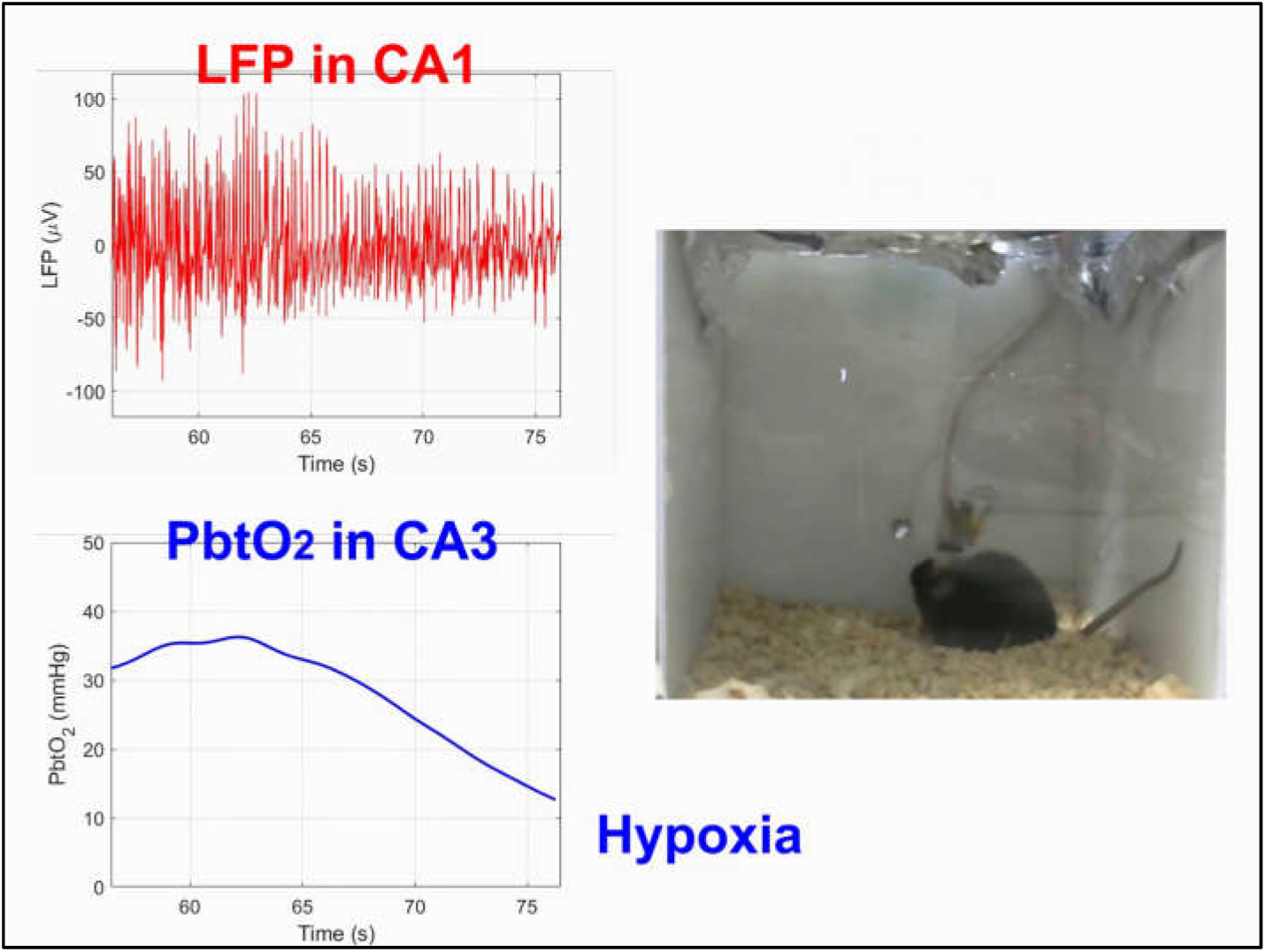
Video (3× speed) showing simultaneously recorded PbtO_2_ in ipsilateral CA3 and LFP signals in CA1, in a freely moving mouse, during afterdischarge induced by electrical stimulation in the CA1.

**Movie S3.**
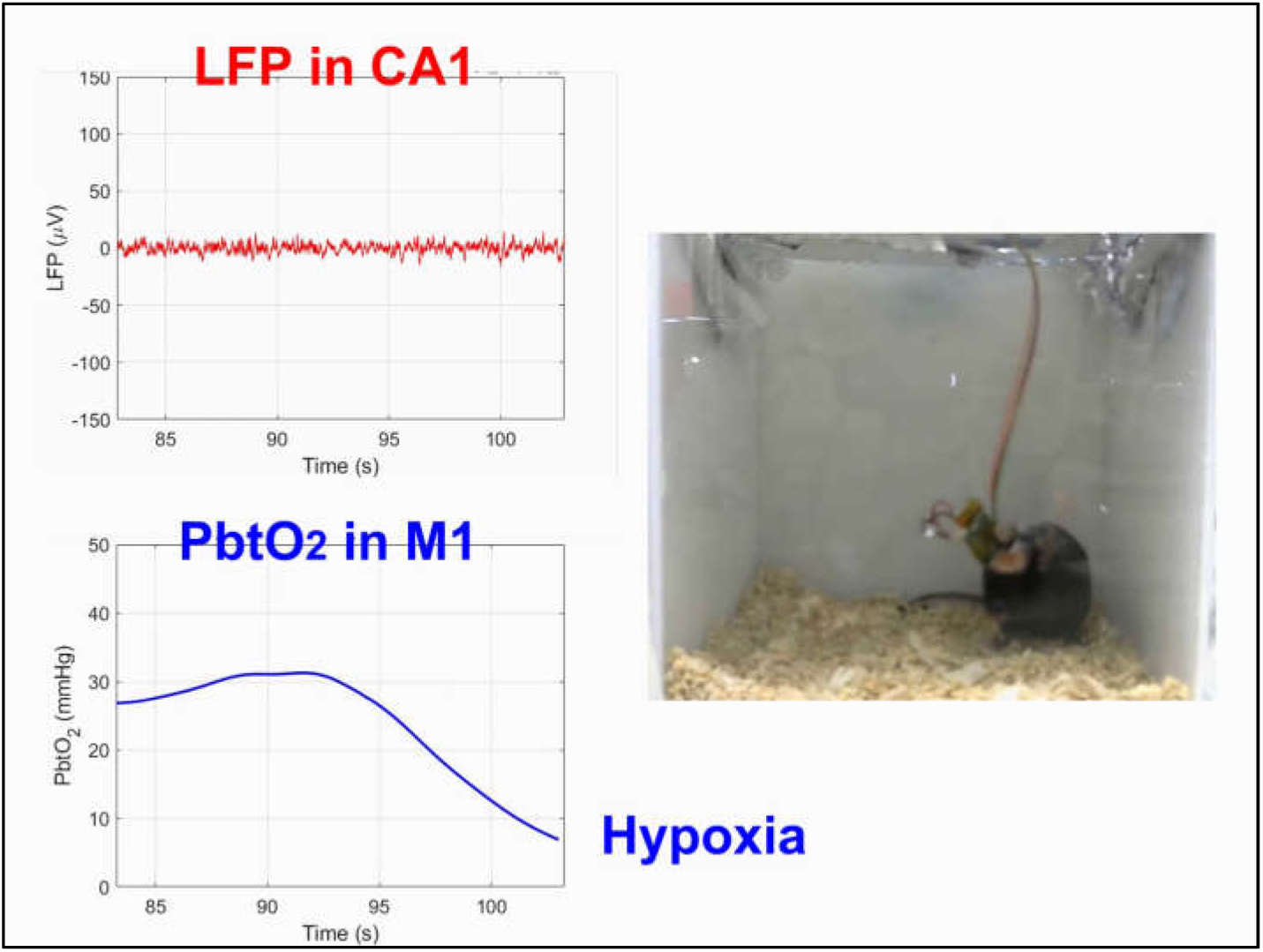
Video (3× speed) showing simultaneously recorded PbtO_2_ in contralateral M1 and LFP signals in CA1, in a freely moving mouse, during afterdischarge induced by electrical stimulation in the CA1.

